# Predator avoidance promotes inter-bacterial symbiosis with myxobacteria in polymicrobial communities

**DOI:** 10.64898/2026.02.12.705600

**Authors:** Shailaja Khanal, Sheila Walsh, Nawal Shehata, Andrew Ahearne, Daniel Belin, Britney Larson, Benjamin Tabor, Daniel Wall, Cole Stevens

## Abstract

Myxobacteria are predatory soil bacteria with the largest known bacterial genomes, rich in biosynthetic gene clusters for specialized metabolites. Despite their ecological importance as potential keystone taxa in soil food webs, there is a disconnect between laboratory-isolated myxobacteria and abundant Myxococcota detected in environmental metagenomic studies. Here, we report the isolation and characterization of stable myxobacterial swarm consortia from rhizospheric soil, consisting of myxobacteria associated with novel *Microvirga* species. Using metagenomic sequencing, we assembled metagenome-assembled genomes (MAGs) for four consortia, revealing phylogenetically distinct yet stably associated bacterial partnerships. Comparative genomics identified evidence of horizontal gene transfer, including acyl-homoserine lactone (AHL) synthases and ankyrin repeat (ANKYR) proteins shared between consortium members, and genome-scale metabolic modeling predicted complementary auxotrophies. Remarkably, time-lapse microscopy revealed that *Archangium* exhibited markedly reduced predation toward its *Microvirga* companion (0.7% predation rate) compared to non-symbiotic *Myxococcus xanthus* (14.9% predation rate), while maintaining robust predatory capacity against *Escherichia coli* prey. These findings indicate that predation avoidance and metabolic complementarity can drive stable inter-bacterial symbiosis in predatory myxobacterial communities, providing foundational insights into previously overlooked myxobacterial partnerships that may be prevalent in natural soil ecosystems.

## Introduction

Members of the phylum Myxococcota, colloquially referred to as myxobacteria, demonstrate traits atypical of bacteria such as “wolf-pack” predatory swarming to acquire macromolecular nutrients from lysed prey and contact-dependent recognition of both kin and prey (1–5). Nearly all myxobacteria from the class Myxococcia are considered generalist predators, where prey includes bacteria, fungi, and oomycetes. Interestingly, their predatory capacity appears to be shaped more by environmental conditions and microbial community composition than phylogeny (6–14). Consistent with ecological impact, recent evidence of bacterial nutrient cycling processes, independent of eukaryotic micropredators, highlighted myxobacteria as potential keystone taxa in the soil food web (15). At the genomic scale, myxobacteria further distinguish themselves by maintaining the largest known bacterial genomes, rich in biosynthetic gene clusters (BGCs), which encode specialized metabolites (16–19). Deemed “gifted” for their potential to produce biologically active metabolites, myxobacteria are targeted for genome mining for discovery of natural products, which have expanded efforts to isolate novel myxobacteria, which have resulted in the discovery of over 40 novel species (20–22). These discoveries include representatives from eleven newly described genera within the last decade. However, the vast majority of laboratory and type strains are minimally present in environmental metagenomic data, and there is a disconnect between myxobacteria with sequenced genomes and abundant Myxococcota from ecological, metagenomic analysis of soil (23–25).

The observation that myxobacteria select for diverse prey during predation may provide insight into this discrepancy with metagenomic sampling (26). Results from controlled predator-prey experiments have repeatedly demonstrated selection of prey phenotypes that influence mucoidy, metabolism, cofactor access, and growth (27–31). Variable prey ranges of myxobacterial isolates suggest these trophic interactions likely scale to environmental conditions in soil and influence microbial community structure (8, 24). We hypothesize that predator-prey coevolution may result in symbiotic relationships between myxobacteria and prey that avoid predation. Predation-resistant neighbors in polymicrobial communities could benefit from shared goods released during myxobacterial lysis of prey populations susceptible to predation. Although laboratory predator-prey experiments have resulted in predation resistant prey, the rapid response and phenotypes observed could also be attributed to general stress responses (28, 32–35). Unfortunately, time and resource constraints limit the likelihood of observing selection of prey resistance and subsequent symbiosis with controlled predator-prey experiments. Similarly, metagenomic investigation of natural polymicrobial communities have provided corollary evidence of associations between myxobacteria and non-myxobacteria, but these analyses cannot elucidate specific symbiotic relationships involving myxobacteria (13, 14, 25). Ultimately, myxobacteria-inclusive consortia isolated from the environment, which are stable and amenable to repeated experimental conditions, are the ideal model to study microbial community structure and possible symbiosis.

Twenty-nine years ago, Jacobi *et al*. reported five isolates of the myxobacterium *Chondromyces crocatus* that were associated with a “companion” bacterium, *Candidatus comitans* (36, 37). Inferring from 16S rRNA sequence homology and observed production of sphingolipids, *Candidatus comitans* was determined to be closely related to members of the genus *Sphingobacterium* and was unable to survive passages as a monoculture removed from co-culture conditions with *C. crocatus*. These myxobacteria-companion pairings would be ideal models for comparative genomic analyses and subsequent, controlled experiments to explore symbiotic traits. However, their discovery predated routine genomics, and no further instances of a naturally occurring myxobacterial consortia have been reported. During our efforts to isolate myxobacteria from soil, we occasionally encounter isolates that we cannot obtain as monocultures, despite repeated passages, and we have also previously reported a contaminant or perhaps companion *Aneurinibacillus* sp. capable of surviving co-culture conditions with the myxobacterium *Archangium violaceum* (38). Searching for myxobacteria-inclusive consortia, similar to the discoveries of Jacobi *et al.*, we sequenced four xenic isolates from rhizospheric soil that were obtained using prey-baiting methodology, but these isolates were notably recalcitrant to yielding myxobacterial monocultures. Mirroring observations from Jacobi *et al.*, these natural swarm consortia grow on solid media as xenic, circular swarms. Herein we report the resulting identities of consortia members, comparative metagenomic findings, and show a companion is specifically resistant to predation by a myxobacteria swarm consortia.

## Results

### Swarm consortia isolated from rhizospheric soil are stable and amenable to laboratory conditions

We sought to isolate swarm consortia using a standard prey-baiting approach that we have previously employed to isolate myxobacteria from rhizospheric soil (20, 39). Our isolation process involves nutrient-free, minimal media supplemented with live *Escherichia coli* as bait; prey-baiting media is inoculated with soil and monitored for the appearance of visible swarms indicative of myxobacterial growth. Observed swarms were assumed to lyse and obtain nutrition from *E. coli* cells. Serial passages of swarms with prey-based minimal media typically yield monocultures of myxobacterial isolates. During this process, three samples WIMLSP1, FLWO, and DLMAZ produced xenic swarms that were recalcitrant to monocultures after repeat passages. Despite numerous passages (*vide infra* in Methods), we were unable to obtain myxobacterial monocultures from WIMLSP1, FLWO, and DLMAZ consortia.

Next, we sought to reisolate WIMLSP1 from its source soil to determine the stability of the candidate swarm consortia and to help rule out the possibility we serendipitously isolated two microbes with similar cultivation conditions. Using the same isolation conditions for WIMLSP1, we were able to isolate a fourth xenic swarm consortia, WIMLSP2. Scanning electron microscopy (SEM) of swarm consortia revealed the presence of cells with two distinct sizes in WIMLSP1 and WIMLSP2 and to a lesser extent FLWO (Figure 1). Larger cells with sizes ranging from 7-9 µM were assumed to be myxobacteria, and smaller cells (1-1.5 µM) were considered candidate companion bacteria. No larger myxobacterial cells were apparent in SEM images of DLMAZ. Subsequent cultivation revealed all four swarm consortia grew on solid and liquid media used for culturing myxobacteria (CTT, CTTYE, CY/H, and VY/2) and endured maintenance as freezer stocks.

**Figure 1:**
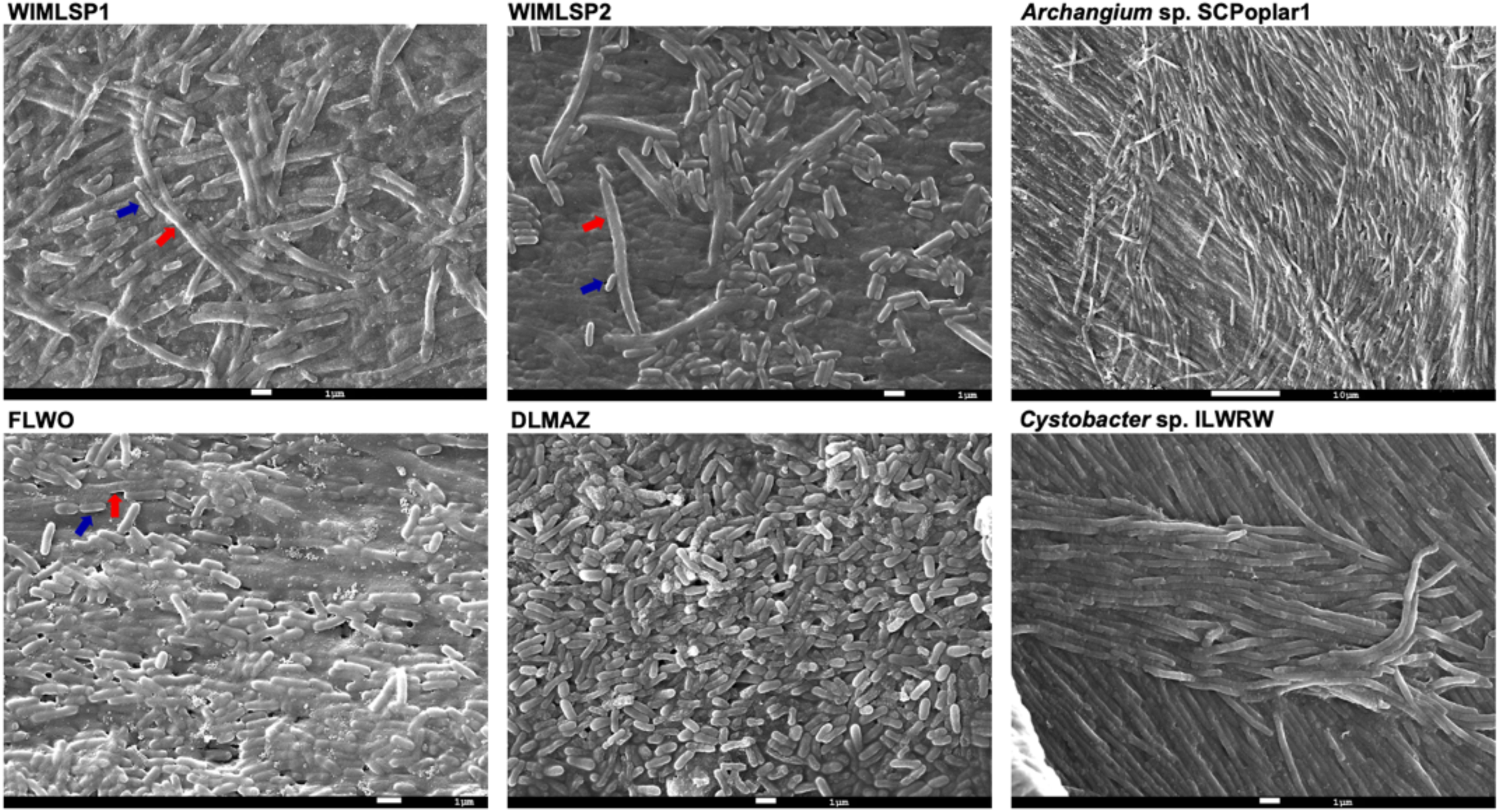
SEM images from swarm consortia WIMLSP1, WIMLSP2, FLWO, DLMAZ and monoculture representatives *Archangium* sp. SCPoplar1 and *Cystobacter* sp. ILWRW for comparison. Representative myxobacterial and microvirgal cells in swarm consortia are labelled with red and blue arrows, respectively.

### Metagenomic analysis reveals association between myxobacteria and *Microvirga* spp

Using metagenomic sequencing, we sought to identify members of each swarm consortia. High molecular weight (HMW) metagenomic DNA was prepared from each consortia and sequenced with long-read nanopore sequencing. Two metagenome assembled genomes (MAGs) were present in each sequenced swarm consortia (Table 1). Consensus data from 16S rRNA comparison, digital DNA-DNA hybridization (dDDH), and average nucleotide identity (ANI) analysis indicated the presence of one myxobacterium and one *Microvirga* spp. in all four consortia. Coverage differences between MAGs from each metagenome assembly were noted as potential differences in abundances between consortia members at the time of HMW DNA isolation. Although fold coverage values for MAGs are not direct measurements of abundance, the differences between consortia are notable with a coverage ratio of 3.75:1 *Archangium* to *Microvirga* in WIMLSP1 and WIMLSP2 and coverage ratios of 5.5:1 and 4.2:1 *Microvirga* to *Archangium*/*Cystobacter* in FLWO and DLMAZ, respectively. Higher abundances of *Microvirga* in FLWO and DLMAZ may explain the relative difficulty we experienced trying to capture images of myxobacterial cells from these consortia using SEM.

**Table 1:** MAG assembly and taxonomy data for sequenced consortia strains.

|  | size<br>(Mb) | coverage | contigs | coding<br>sequences | GC<br>content | 16S rRNA | ANI |
| --- | --- | --- | --- | --- | --- | --- | --- |
| ***Archangium***<br>**sp. WIMLSP1** | 12.98 | 150x | 1 | 10,867 | 68.8% | 99.8%<br>(*Archangium*<br>*gephyra*) | 97.2% (*A.*<br>*violaceum*) |
| ***Microvirga***<br>**sp. WIMLSP1** | 3.9 | 40x | 1 | 3,840 | 61.2% | 99.3%<br>(*Microvirga* sp.<br>GI_Sw3_5a_4) | 84.4%<br>(*Microvirga*<br>*solisilvae*) |
| ***Archangium***<br>**sp. WIMLSP2** | 12.93 | 150x | 1 | 10,737 | 68.7% | 99.8% (*A.*<br>*gephyra*) | 97.2% (*A.*<br>*violaceum*) |
| ***Microvirga***<br>**sp. WIMLSP2** | 4.29 | 40x | 1 | 3,840 | 61.2% | 99.3%<br>(*Microvirga* sp.<br>GI_Sw3_5a_4) | 82% (*M.*<br>*solisilvae*) |
| ***Archangium***<br>**sp. FLWO** | 13.96 | 40x | 1 | 11,642 | 68.1% | 99.8%<br>(*Archangium*<br>*lansingense*) | 96.4% (*A.*<br>*lansingense*) |
| ***Microvirga***<br>**sp. FLWO** | 3.82 | 220x | 1 | 3,658 | 60.9% | 99.1%<br>(*Microvirga* sp.<br>GI_Sw3_5a_4) | 84.5% (*M.*<br>*solisilvae*) |
| ***Cystobacter***<br>**sp. DLMAZ** | 12.01 | 84x | 1 | 10,528 | 68.5% | 99.9%<br>(*Cystobacter*<br>*fuscus*) | 99.3%<br>(*Cystobacter*<br>*ferrugineus*) |
| ***Microvirga***<br>**sp. DLMAZ** | 4.13 | 351x | 1 | 4,001 | 61.4% | 99% (*Microvirga*<br>sp. MB12) | 81.9% (*M.*<br>*solisilvae*) |

Utilizing ANI and dDDH values according to established methods for taxonomic placement (40, 41), we determined *Archangium* present in WIMLSP1 and WIMLSP2 share 99.2% ANI, and both are sub-species of the type strain *Archangium violaceum* with ∼97.2% ANI. *Archangium* sp. FLWO is a sub-species of the type strain *Archangium lansingense* (96.4% ANI), and *Cystobacter* sp. DLMAZ shares 99.3% ANI with the sequenced strain *Cystobacter ferrugineus* and 93.4% ANI with the type strain *Cystobacter fuscus*. Notably, none of the myxobacterial MAGs met the established thresholds to be considered novel species (Figure 2a and b).

**Figure 2:**
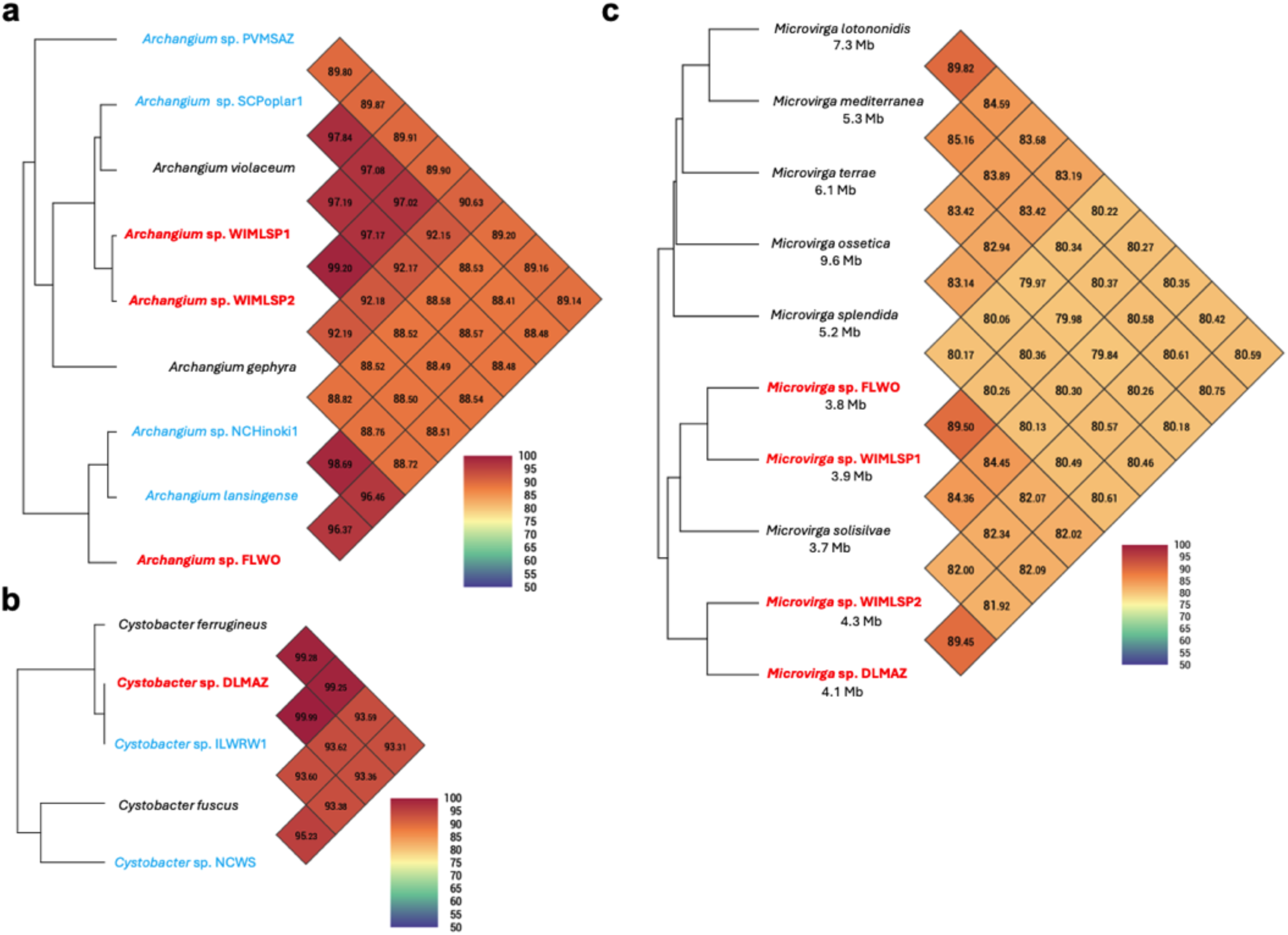
Average nucleotide identity heatmaps for *Archangium* (a), *Cystobacter* (b), and *Microvirga* (c). MAGs from swarm consortia (red) compared with type strain representatives (black) and environmental isolates (blue). Genome sizes for *Microvirga* are included to denote the differences in genome sizes between monoculture and swarm consortia *Microvirga*. Heatmaps generated using OAT (88).

In contrast, all companion *Microvirga* are candidate novel species with <85% ANI shared between sequenced MAGs and the most similar type strain, *Microvirga solisilvae* (42). These ANI values are well below the established threshold (<95% ANI) to be considered a novel species. *Microvirga* from each swarm consortia share <90% ANI and appear to be distinct species, including *Microvirga* sp. WIMLSP1 and *Microvirga* sp. WIMLSP2. From these results, we determined that WIMLSP1 and WIMLSP2 are different swarm consortia with highly related *Archangium* sp. (99.2% ANI) and much less related *Microvirga* sp. (82.3% ANI). Other than the closest relative type strain species, *Microvirga solisilvae*, swarm consortia-associated *Microvirga* have much smaller genomes (∼3.7-4 Mb) than sequenced, monocultured *Microvirga* (5.2-9.6 Mb) (Figure 2c). From these observations we hypothesize symbiosis in swarm communities may have contributed to *Microvirga* gene loss over time resulting in genome reduction.

### Comparative genomics indicate horizontal gene transfer in swarm communities

With MAG sequence data for each consortia strain in hand, we sought additional attributes of swarm communities that would be indicative of symbiosis. The Joint Genome Institute (JGI) Integrated Microbial Genomes and Microbiomes (IMG/MER) database was used to analyze the phylogenetic distribution of genes from consortia metagenomes to assess potential horizontal gene transfer within swarm consortia (43, 44). Using this approach, LuxI-like acyl-homoserine lactone (AHL) synthases homologous to *Microvirga* were identified in *Archangium* MAGs from WIMLSP1 and WIMLSP2 (Table 2) (45, 46). Both LuxI homologs share >70% amino acid identity with an acyl-homoserine lactone synthase from *Microvirga* sp. 2TAF3. LuxI-like AHL synthases from *A.* WIMLSP1 and *A.* WIMLSP2 were also highly homologous to the only known myxobacterial AHL synthases previously discovered from *A. gephyra* (>90 identity) and *Vitiosangium* sp. GDMCC 1.1324 (>70% identity) (47). LuxI homologs were also present in *Microvirga* MAGs from WIMLSP1 and WIMLSP2. Foldseek analysis of cognate LuxI structures from each swarm consortia reveals structural similarity (Figure 3a and 3b), and phylogenetic analysis corroborates a shared evolutionary history between *Archangium* and *Microvirga* AHL synthases (Figure 3c) (48, 49). These results suggest horizontal transfer of an AHL synthase gene from *Microvirga* may account for the AHL synthases present in *A.* WIMLSP1, *A.* WIMLSP2, *A. gephyra*, and *V.* GDMCC 1.1324.

**Figure 3:**
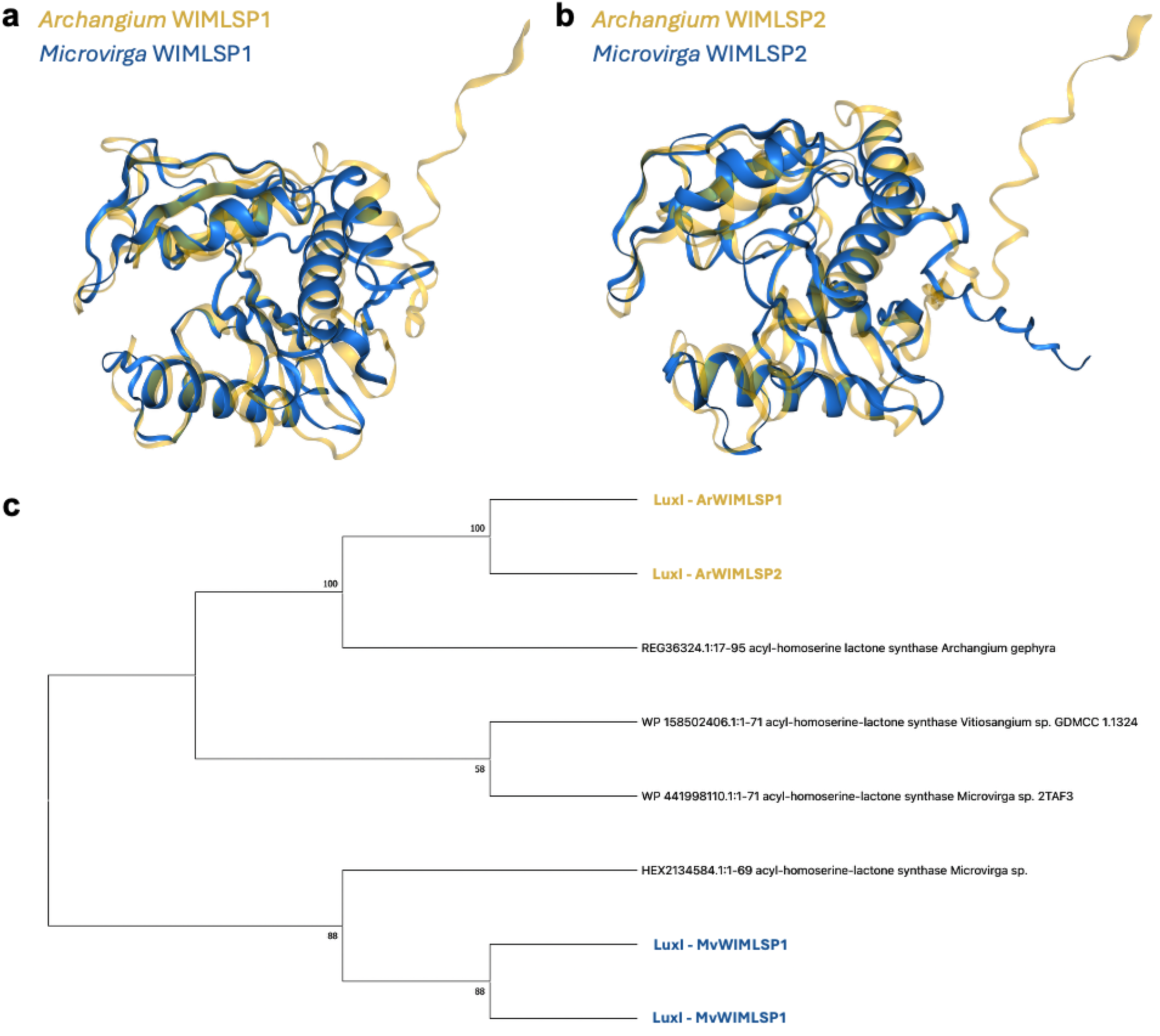
Alignment of structural models for AHL synthases present in each member of WIMLSP1 (a) and WIMLSP2 (b) swarm consortia. Evolutionary analysis of myxobacterial and microvirgal AHL synthases (c) from a bootstrap consensus tree inferred from 300 replicates. Values shown next to branches are the percentage of replicated trees with the associated taxa clustered together. Structural models of AHL synthases were generated using the AlphaFold Server and aligned with FoldMason MSA (48, 49). Evolutionary history was inferred using the Maximum Likelihood method and JTT matrix-based model in MEGA X (89, 91–93).

**Table 2:**
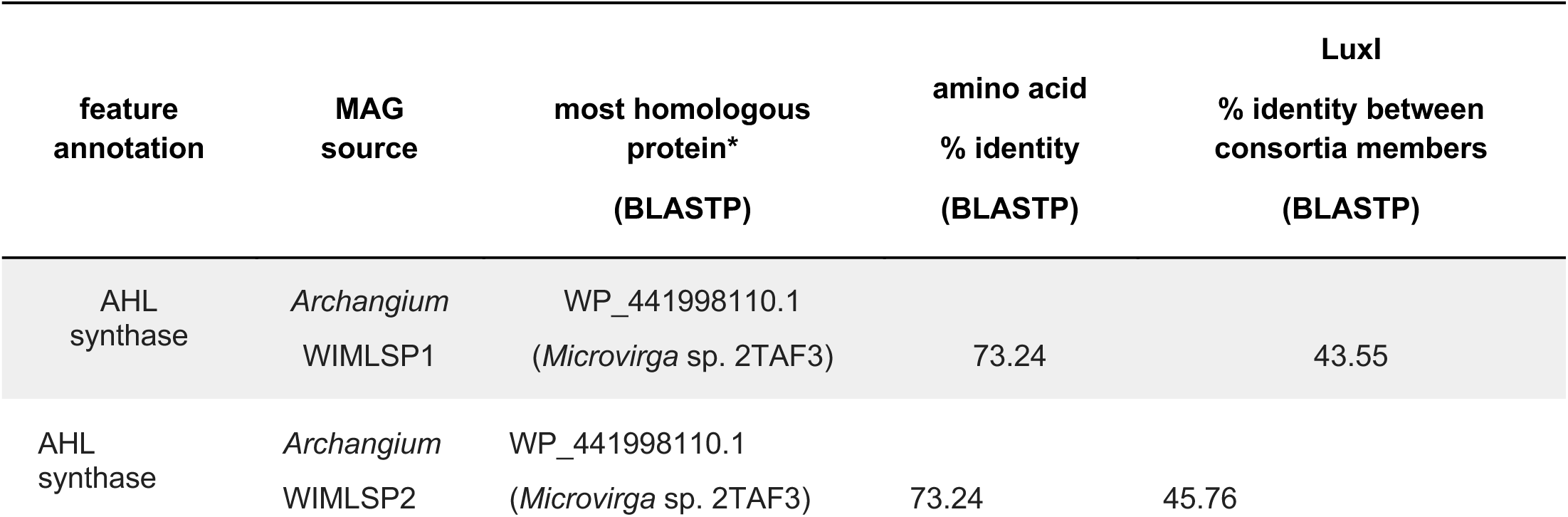
Annotated LuxI homologs from *Archangium* MAGs that share homology with *Microvirga*. *Excluding LuxI homologous from Myxococcota.

Proteins with homology to myxobacterial proteins were found in all *Microvirga* MAGs. All *Microvirga* MAGs include encoded proteins homologous with *Archangium* proteins (Table 3), including *Microvirga* sp. DLMAZ which has no gene products that share homology with sequenced *Cystobacter*. Subsequent analysis revealed homologous ankyrin repeat domain-containing (ANKYR) proteins present in all members of WIMLSP1 and WIMLSP2 swarm consortia (Table 3; bolded rows). A sequence similarity network (SSN) generated from the shared ANKYR proteins using the Enzyme Function Initiative Enzyme Similarity Tool (EFI-EST) identified 15 homologous entries in the UniProt database (Supplemental Figure S5) (50, 51). Homologous ANKYR proteins in the generated SSN are exclusively present in Archangiaceae within the phylum Myxococcota. Although EFI-EST analysis identified ANKYR proteins variably present in eight genera from the phylum Pseudomonadati, none were from the genus *Microvirga*. Comparative genomic analysis of the 157 sequenced *Microvirga* available at the National Center for Biotechnology and Information (NCBI) genome database revealed the presence of ANKYR protein homologs in just two additional *Microvirga*, *Microvirga solisilvae* and *Microvirga* sp. ACRRW. Comparing the spatial organization of ANKYR encoding genes for *M. solisilvae*, *M.* sp. ACRRW, *M.* WIMLSP1, and *M.* WIMLSP2, we observed a neighboring XerC tyrosine recombinase in each genome (Supplemental Figure S7). XerC-dependent phage integration has been previously reported (52–54), and proximity to all ANKYR-encoding genes present in sequenced *Microvirga* buttresses the likelihood of horizontal acquisition. ANKYR proteins from WIMLSP1 and WIMLSP2 also share high nucleotide identities with >65% coverage values (Table 3; Supplemental Figure S6), modelled structural homology (Figure 4a and 4b), and evolutionary histories supported by phylogenetic analysis (Figure 4c). Taken together, our data suggests horizontal transfer of ANKYR proteins from *Archangium* to *Microvirga* in WIMLSP1 and WIMLSP2.

**Figure 4:**
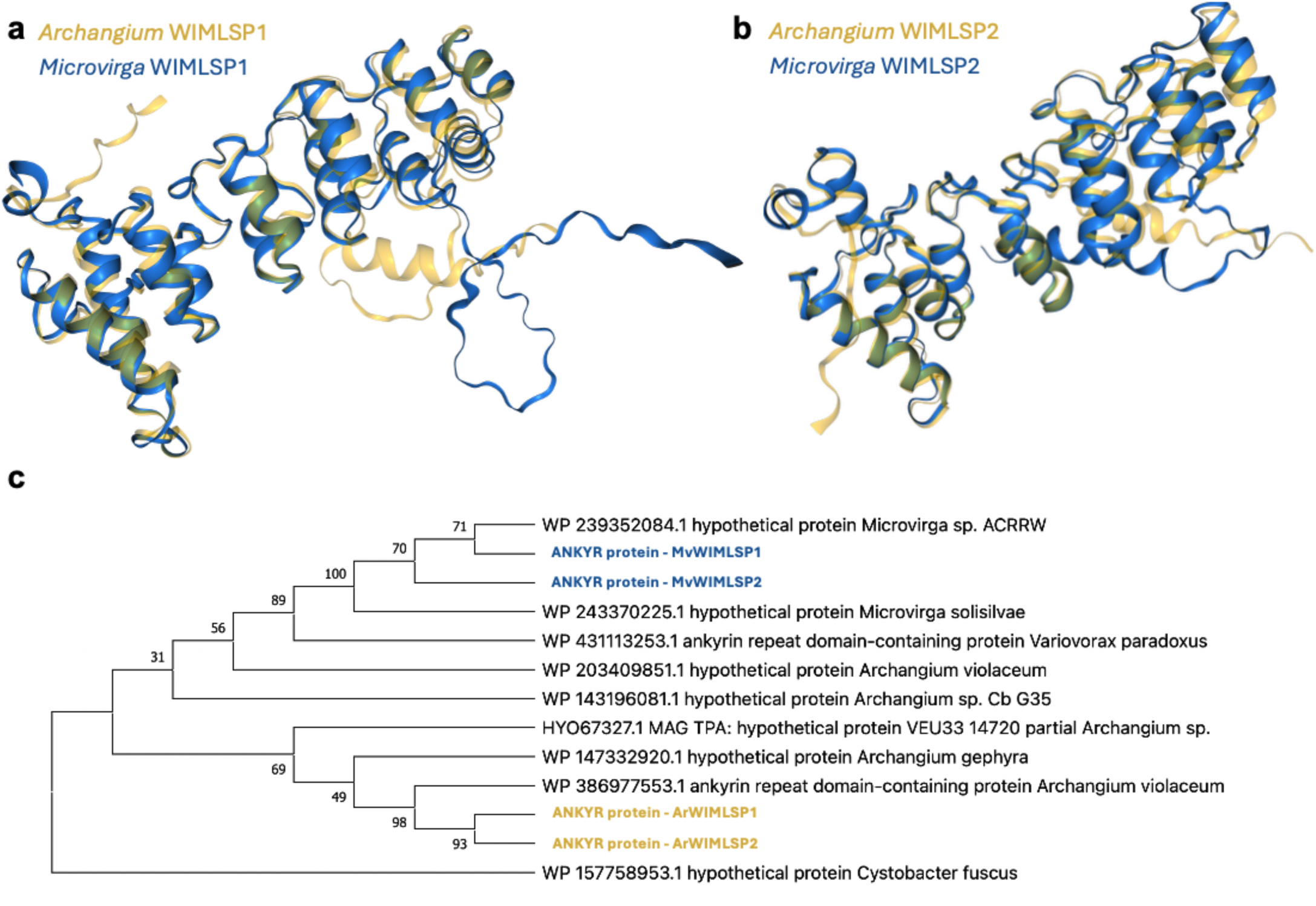
Alignment of structural models for ANKYR proteins present in each member of WIMLSP1 (a) and WIMLSP2 (b) swarm consortia. Evolutionary analysis of myxobacterial and microvirgal ANKYR proteins (c) from a bootstrap consensus tree inferred from 300 replicates. Values shown next to branches are the percentage of replicated trees with the associated taxa clustered together. Structural models of AHL synthases were generated using the AlphaFold Server and aligned with FoldMason MSA (48, 49). Evolutionary history was inferred using the Maximum Likelihood method and Le_Gascuel_2008 model in MEGA X(89, 91–94).

**Table 3:**
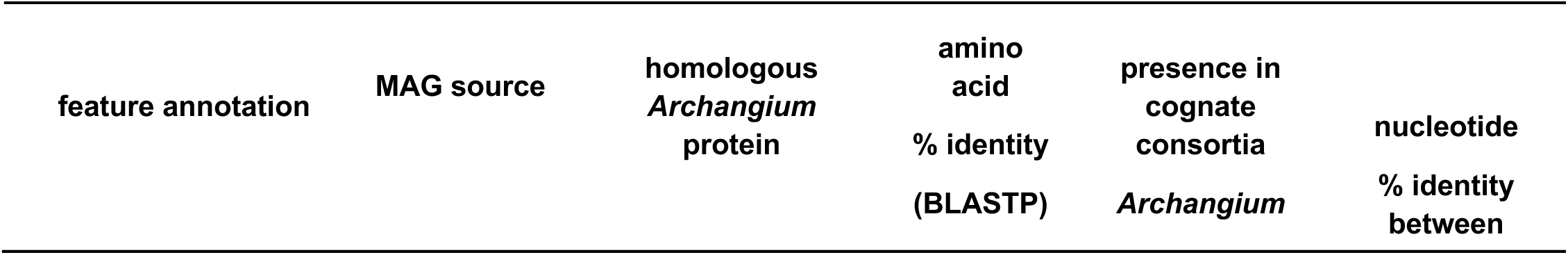
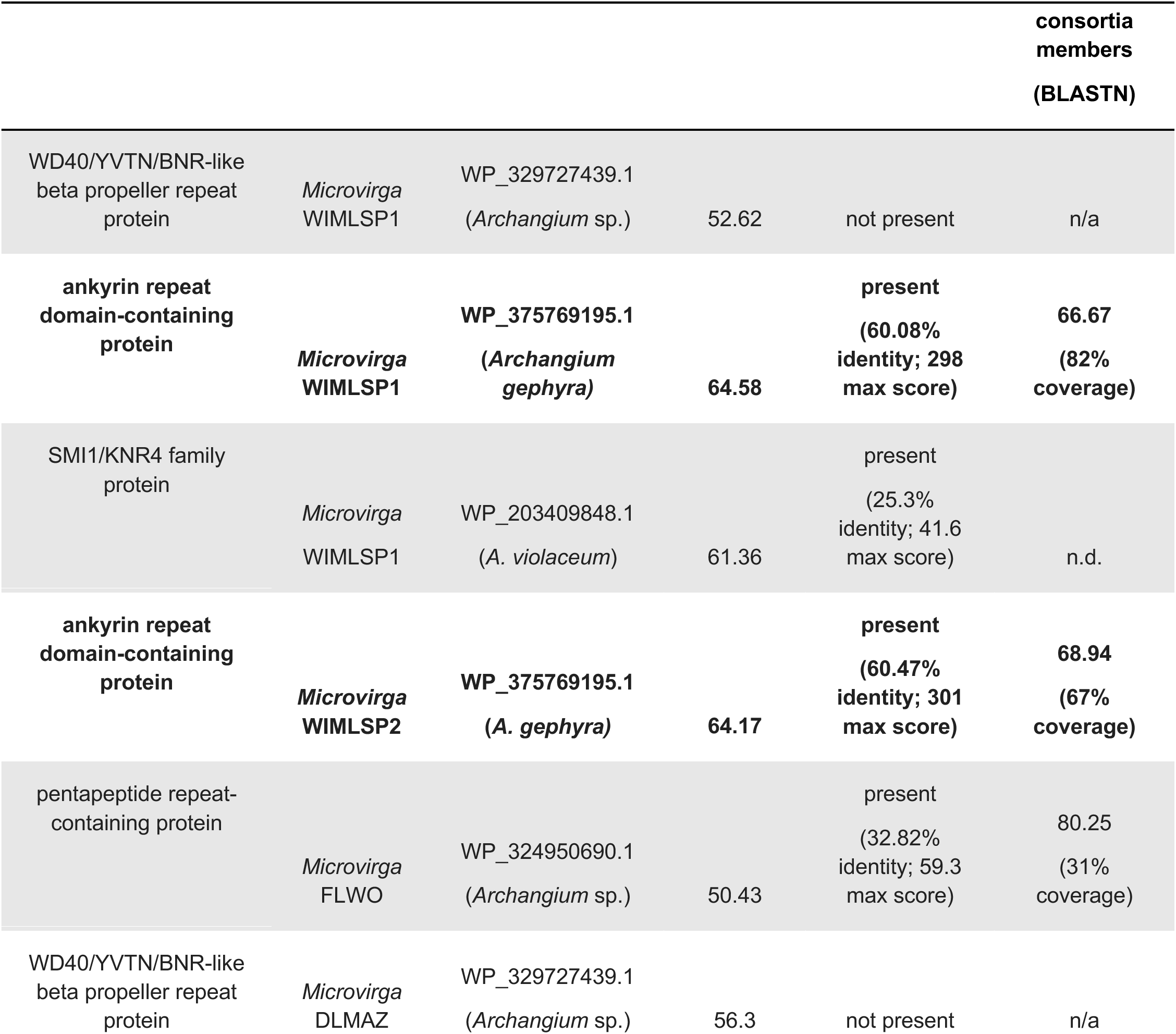
Features from *Microvirga* MAGs that share homology with *Archangium*. Proteins with high homology present in both members of a swarm consortia are bolded.

### Auxotrophies and potential metabolic exchanges in swarm consortia identified by genome-scale metabolic modeling

Metabolic exchanges in consortia that alleviate auxotrophies of members would also indicate a basis for inter-bacterial symbiosis i (52). Genome-scale metabolic models (GEMs) were built for consortia MAGs and related monoculture species for comparison using the ModelSEED2 pipeline and subsequently characterized to predict auxotrophy (Tables 4 and 5)(53–55). All swarm consortia MAGs were secondarily analyzed with GapMind to corroborate predicted amino acid auxotrophies (56). All *Archangium* consortia members are predicted to be branched-chain amino acid (BCAA) auxotrophs as well as L-histidine, folate, and riboflavin auxotrophs (Table 4). Myxobacteria are often BCAA auxotrophs that require acquisition of BCAAs from prey lysates (57). Comparing consortia *Archangium* with sequenced, monocultured *Archangium*, these appear to be somewhat general auxotrophies within the genus. A similar comparison of consortia and monoculture *Cystobacter* revealed no common auxotrophies within the genus, and *C.* DLMAZ was found to only share L-histidine auxotrophy with analyzed *Archangium*. Comparative analysis of consortia metagenomes using assigned KEGG orthologies confirmed the absences of dihydroxy-acid dehydratase *ilvD*, 2-isopropylmalate synthase *leuA*, and citramalate isomerase subunits *leuCD* required for BCAA biosynthesis in *Archangium* from WIMLSP1, WIMLSP2, and FLWO (Supplemental Figure S8) (58–61).

**Table 4:** Auxotrophies (A) and prototrophies (P) of consortia *Archangium* (bolded) and monoculture *Archangium*. *L-isoleucine auxotrophy in *Archangium* was predicted by GapMind analysis but was not observed in metabolic models(56).

|  | L-His | L-Leu | L-Val | L-Ile* | folate | riboflavin |
| --- | --- | --- | --- | --- | --- | --- |
| ***Archangium* sp. WIMLSP1** | **A** | **A** | **A** | **A** | **A** | **A** |
| ***Archangium* sp. WIMLSP2** | **A** | **A** | **A** | **A** | **A** | **A** |
| ***Archangium* sp. FLWO** | **A** | **A** | **A** | **A** | **A** | **A** |
| *Archangium* sp. PVMSAZ | **A** | **A** | **A** | **A** | **A** | **A** |
| *Archangium* sp. SCPoplar1 | **A** | **A** | **A** | **A** | **A** | **A** |
| *Archangium* *lansingense* | P | **A** | **A** | **A** | P | P |
| *Archangium* *gephyra* | P | **A** | **A** | **A** | P | P |
| ***Cystobacter* sp. DLMAZ** | **A** | P | P | P | P | P |
| *Cystobacter* *ferrugineus* | P | P | P | P | P | P |

**Table 5:**
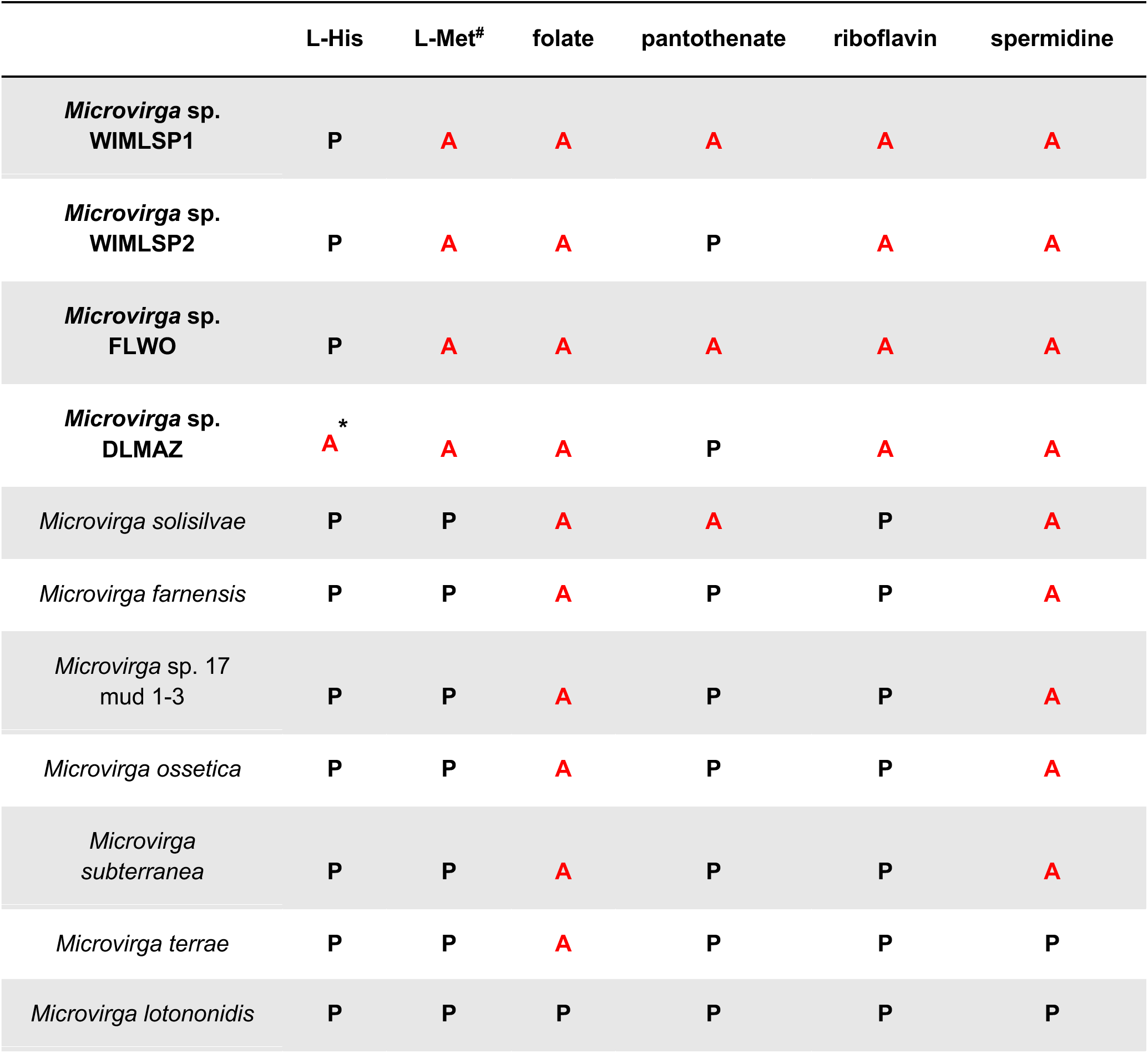
Auxotrophies of consortia *Microvirga* (bolded) and monoculture *Microvirga*. *Auxotrophies predicted by GapMind analysis but not metabolic models. ^#^L-methionine auxotrophy in *Microvirga* was predicted by metabolic models but was not observed in Gapmind analysis(56).

Predicted auxotrophies of consortia-associated and monoculture *Microvirga* were more varied with minimal overlap in auxotrophies between strains (Table 5). All four *Microvirga* swarm consortia were predicted to be L-methionine, folate, and riboflavin auxotrophs from GEM analysis. However, subsequent Gapmind analysis predicted all consortia *Microvirga* to be methionine prototrophs with medium confidence. None of the analyzed monoculture *Microvirga* were predicted to be L-methionine auxotrophs by either GEM or Gapmind analysis. Methionine is considered a biosynthetically costly amino acid, and methionine auxotrophy has been found to promote metabolic exchanges between symbionts (62). Using KEGG orthologies to assess predicted L-methionine auxotrophy, we confirmed the absence of the homoserine *O*-succinyltransferase *metA* required for L-methionine biosynthesis via L-cystathionine in all four consortia-associated *Microvirga*(63). Alternatively, all four *Microvirga* were found to have a complete sulfhydrylation pathway that utilizes hydrogen sulfide for L-methionine production (64). Discrepancies between GEM analysis and Gapmind analysis are likely due to the intrinsic homology between homoserine *O-*succinyltransferase *metA* from the L-cystathionine-dependent methionine biosynthetic pathway and *metX* from the sulfhydration pathway. Intrigued by this observation, we next verified the absence of an assimilatory sulfate reduction pathway in all four *Microvirga* to exclude potential L-methionine prototrophy via direct sulfhydrylation of *O*-acetyl-L-homoserine to L-homocysteine(65). These observations were mirrored in consortia-associated *Archangium* with no observed *metA* orthologs. However, complete direct sulfhydrylation L-methionine biosynthetic pathways are present in all four *Archangium*.

Identification of potential nutrient exchanges in swarm consortia suggests consortia stability and support inter-bacterial symbiosis. Exchange of BCAAs from *Microvirga* to *Archangium* provides a straightforward opportunity for cross-feeding in swarm consortia (Figure 5). Accumulation of BCAAs and various intermediate metabolites has been shown to inhibit BCAA biosynthesis (66), and BCAA consumption by *Archangium* would promote BCAA biosynthesis in *Microvirga*. The overlap in direct sulfhydrylation L-methionine biosynthetic pathways between consortia members provided an additional potential nutrient exchange. Hydrogen sulfide exchange from producing *Archangium*, via intact assimilatory sulfate reduction pathways, would enable L-methionine biosynthesis from *Microvirga* in swarm consortia (Figure 4). Biosynthesis of L-methionine by consortia *Archangium*, may also provide L-methionine directly to consortia *Microvirga* independent of hydrogen sulfide exchange. These results provide potential nutrient exchanges between members of swarm consortia that would alleviate essential amino acid auxotrophies.

**Figure 5:**
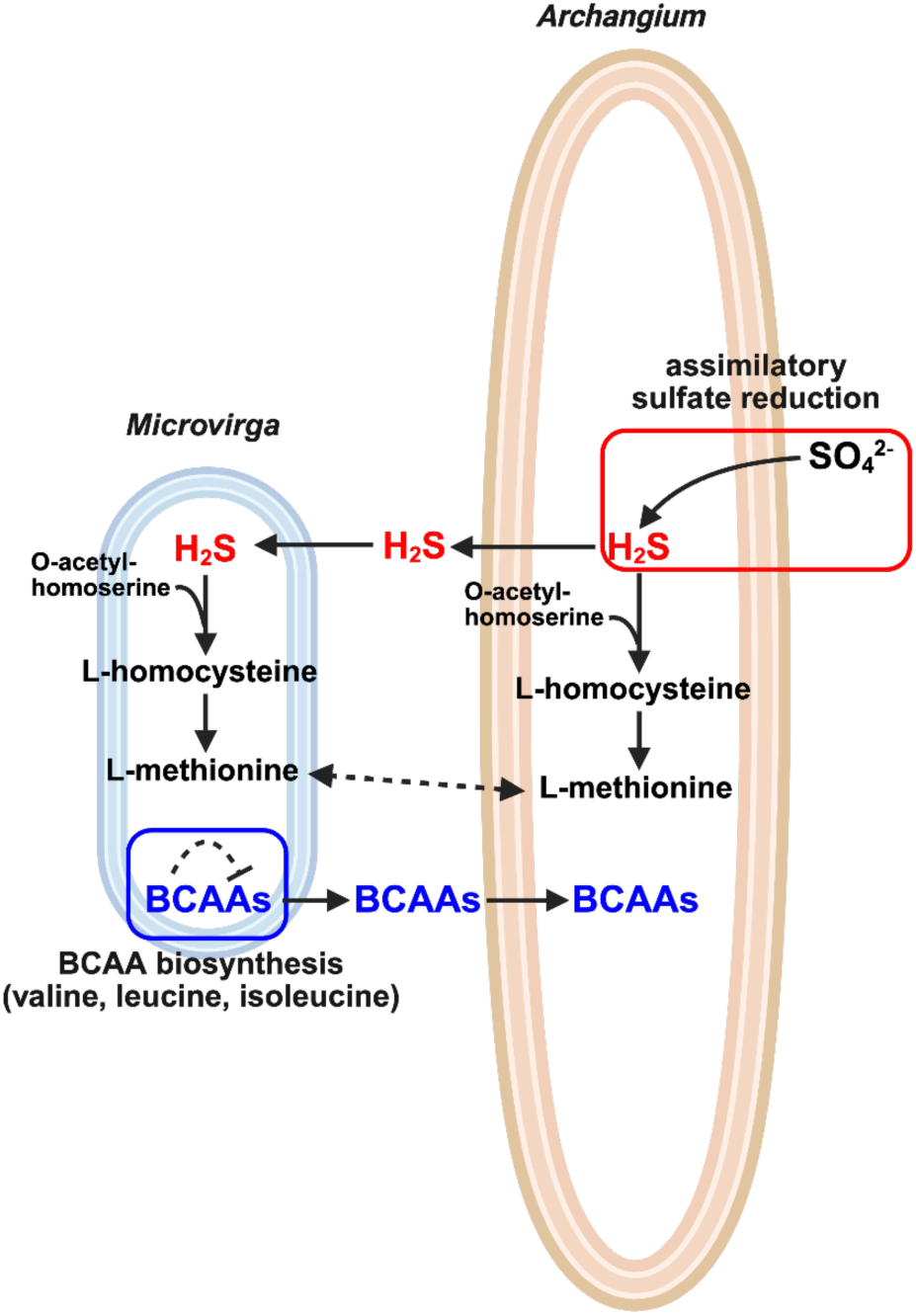
Proposed metabolic exchanges in WIMLSP1 and WIMLSP2 to alleviate L-methionine auxotrophy in *Microvirga* and BCAA auxotrophy in *Archangium*. Image generated in BioRender.

### Intra-consortia predation of *Microvirga* companions was rare

Predatory lifestyles of myxobacteria in the class Myxococcia — including *Archangium —* complicate the possibility of stable symbiosis within swarm consortia, as myxobacteria are known to exhibit exceptional predatory capacities. Predation of Alphaproteobacteria such as *Sinorhizobium meliloti* and *Agrobacterium tumefaciens* has been documented (67, 68). To assess if predation occurs within a naturally occurring consortia, we leveraged distinct cell-size differences between members and selected the WIMLSP2 consortium as the best model consortia due to the favorable ratio of members observed previously by SEM. Time-lapse movies of WIMLSP2 were recorded, and cell-cell interactions between consortia members from WIMLSP2 were enumerated to record the number of times that *M.* WIMLSP2 cells lysed following contact with *A.* WIMLSP2. Of the 1,299 observed cell-cell interactions, only 23 resulted in *Microvirga* lysis, yielding an intra-consortium predation rate of just 2.5% (Figure 6). These results indicate that predation of *Microvirga* companions by *Archangium* was rare within the swarm consortium WIMLSP2, supporting the interpretation that these species coexist stably rather than primarily engaging in predator-prey interaction.

**Figure 6:**
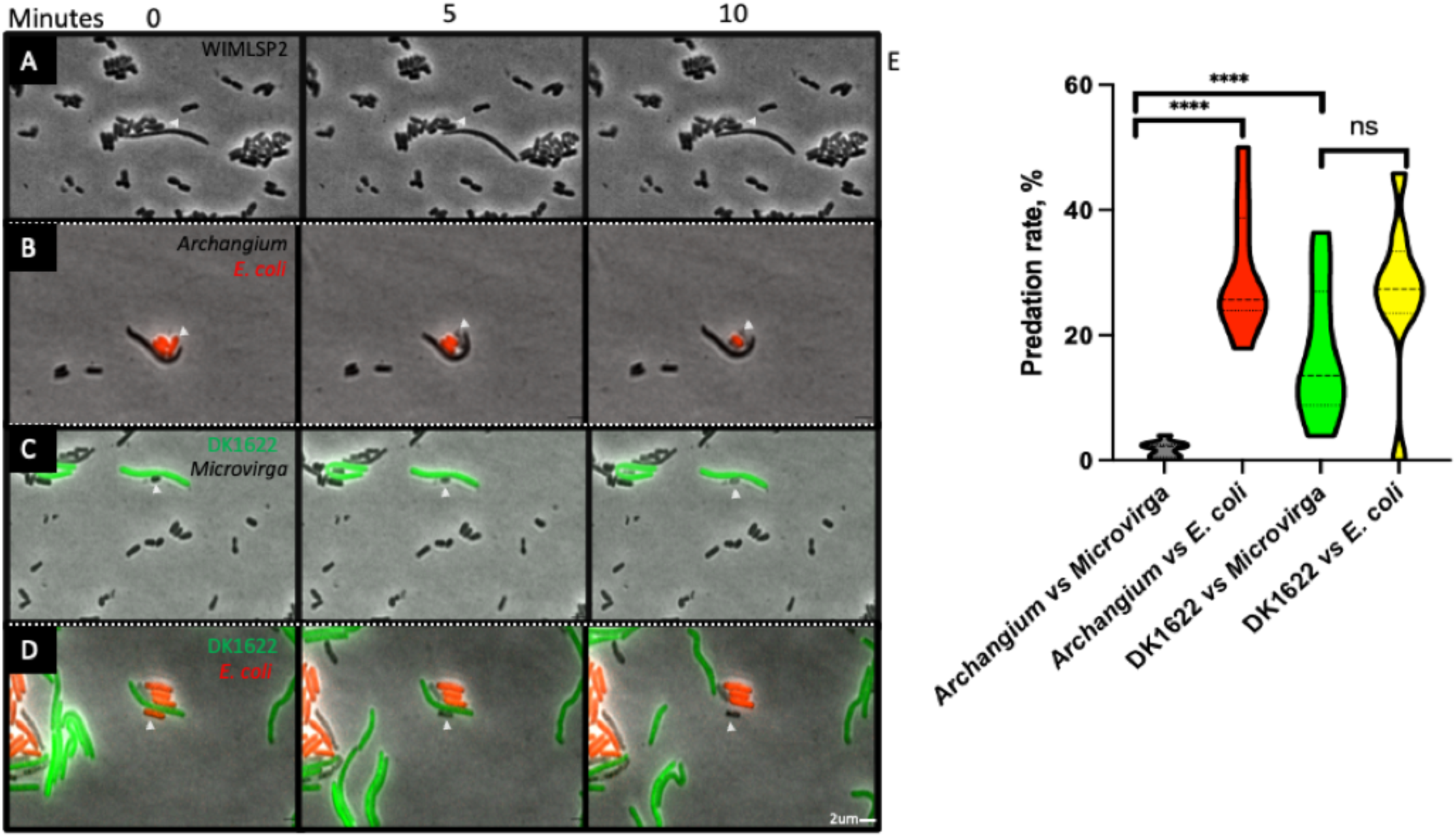
Contact-dependent predation of bacterial prey by myxobacteria. Time-lapse microscopy stills demonstrate predator-prey interactions at 0, 5 and 10 minutes following initial contact (white arrowheads indicate prey cells). (A) *Archangium* WIMLSP2 interacting with *Microvirga* WIMLSP2 cells. (B) *Archangium* WIMLSP2 interacting and killing *E. coli* cells. (C) DK1622 interacting with and killing *Microvirga* WIMLSP2. (D) *M. xanthus* DK1622 interacting with and killing *E. coli*. (E) Violin plots illustrating the distributions of predation rates (%) calculated from time-lapse movies for each predator-prey pairing. Each violin represents the full distribution of predation rates, with the dashed lines indicating the median and dotted lines indicating the interquartile range. Statistical comparisons were performed using a two-tailed Mann-Whitney U test. **** indicated *p*< 0.0001; ns, not significant.

### Confirmed predation of *E. coli* by swarm consortia

We next revisited the predatory capacity of WIMLSP2 using *E. coli* as prey to test whether the low intra-consortium predation rate reflected *bona fide* resistance by *Microvirga* rather than a defect in *Archangium* predation. Although all swarm consortia were originally isolated using prey-baiting with *E. coli*, we sought to enumerate consortia predation with a mCherry-labelled *E. coli* prey. Fluorescence *E. coli* cells enabled us to similarly track *A.* WIMLSP2 contact with *E. coli* and document subsequent lysis events. A total of 139 lysis events were observed after monitoring 502 cell-cell interactions between *A.* WIMLSP2 and introduced *E. coli* prey. The calculated 28% predation rate from these results demonstrate WIMLSP2 capably predates *E. coli* that is significantly different than *Microvirga* lysis rate (Figure 6). For comparison, we performed the same assay using *Myxococcus xanthus* DK1622-GFP and *E. coli-*mCherry and observed 278 lysis events out of 930 total total cell-cell interactions, confirming the expected predatory efficiency of the model myxobacterium *My. xanthus* DK1622 (69). These results show *A.* WIMLSP2 and *My. xanthus* have comparable *E. coli* predation rates (28% and 30%) and suggest *Microvirga* swarm companions may be refractile to myxobacteria predation.

### M. *xanthus* predates *Microvirga* companions

Using the same approach to determine the predation rate of *Microvirga* WIMLSP2 by *My. xanthus* DK1622-GFP, we monitored 1,339 total predator–prey interactions and recorded 199 lysis events. From these results, a predation rate of 15% was calculated (Figure 6). This rate is higher than that observed for predation by *A.* WIMLSP2, indicating that *M.* WIMLSP2 is more susceptible to DK1622-GFP than to its native myxobacterial partner.

## Discussion

Our discovery of myxobacterial swarm consortia corroborates the myxobacterial companionships reported by Jacobi et al. nearly three decades ago (36, 37). Phylogenetic differences between myxobacteria present in each reported swarm consortia, with *Chondromyces* from the subclass Polyangiaa and *Archangium* from the subclass Myxococcia, suggest inter-bacterial symbiosis with myxobacteria may be prevalent in polymicrobial communities. Supporting the potential prevalence of swarm consortia, a recent investigation of biotic interactions in the alpine soil microbiome during seasonal shifts observed positive associations between Alphaproteobacteria and Myxococcota (70). Alphaproteobacterial genomes have extraordinary plasticity associated with facultative, intracellular, and free-living lifestyles reported from the class, and horizontal gene transfer occurs commonly in Alphaproteobacteria (71). Genetic adaptability is reflected in the substantial differences in alphaproteobacterial genome sizes, ranging from 1-9 Mb. The lifestyle differences of *Microvirga* exemplify the genetic variability of Alphaproteobacteria (72). The genus currently has 27 species with valid descriptions that have been isolated from a range of environments other than soil including the atmosphere, hot springs, deep sea thermal aquifers, plant root nodules, metal industry waste, and human stool. Described *Microvirga* strains also include plant and human symbiotes demonstrating precedence for symbiosis in the genus (72). *Microvirga* from swarm consortia appear to be novel species, and are the first *Microvirga* found inter-bacterial symbionts, and the second reported “companions” of predatory myxobacteria. Noted metabolic exchanges and horizontally transferred AHL synthases and ANKYR gene products are indications of symbiosis.

Although exogenous AHLs have been shown to elicit responses from myxobacteria, the underlying mechanisms remain unknown, and myxobacteria are not known to participate in AHL-mediated quorum sensing (73, 74). There are no documented instances of a complete LuxR/LuxI quorum sensing system in a myxobacterium. The previously mentioned functional AHL synthases from *A. gephyra* and *V.* GDMCC 1.1324 are orphaned without a paired LuxR receptor, and AHL production has not been directly observed from either myxobacterium (47). Each myxobacterial LuxI-homolog required heterologous expression in an *E. coli* host to confirm functionality and biosynthesis of AHLs. Our data suggests AHL synthases from *A.* WIMLSP1 and *A.* WIMLSP2 are highly similar to the *A. gephyra* and *V.* GDMCC 1.1324 AHL synthases. Phylogenetic analysis also supports a potential shared evolutionary history between AHLs synthases from myxobacteria and *Microvirga*. Horizontal acquisition of biosynthetic genes has been well-documented in myxobacteria (20, 75), and we are intrigued to further explore quorum sensing in swarm consortia.

Interestingly, ANKYR proteins have been implicated in host immune avoidance in a variety of symbiotic relationships. As examples, ANKYR proteins VAPYRIN and IGN1 are required for host accommodation in fungal- and bacterial-plant symbioses (76, 77), and ANKYR genes of the intracellular *Wolbachia* symbionts rapidly evolve and have been suggested to contribute to *Wolbachia*-*Drosophila* symbiosis (78–80). We are excited to develop genetic approaches suitable for swarm consortia engineering, which will enable further investigation of ANKYR proteins shared by members of WIMLSP1 and WIMLSP2 to explore any potential contributions to myxobacterial symbiosis.

The presence of distinguishable microvirgal symbionts, *M.* WIMLSP1 and *M.* WIMLSP2, associated with practically the same strain of *Archangium* from the same soil sample, suggest symbiosis in swarm consortia is not exclusive and may be transient. Notably, we previously isolated monocultures of *Archangium* sp. SCPoplar1, *Archangium lansingense*, and *Cystobacter* sp. ILWRW that are nearly identical (97% and 99% ANI respectively) to myxobacteria from swarm consortia (20, 75). We hypothesize *Microvirga* that have adapted to avoid predation to certain myxobacterial, establish transient symbiotic relationships with them in polymicrobial communities and benefit from nutrients released by myxobacterial lysis of prey. *Microvirga* have been noted to be common “contaminates” that complicate or prevent successful isolation of axenic myxobacteria from soil (81). Our observations that support this hypothesis include genes phylogenetically associated with a variety of myxobacteria present in genomes of consortia *Microvirga* (Supplemental Table S8) and the genetic variability of consortia-associated *Microvirga*. For example, the BNR-like beta propeller repeat proteins from *M.* WIMLSP1 and *M.* DLMAZ with homologs in *Archangium* (Table 3) only share 55% amino acid identity and may have been acquired from previous myxobacterial symbionts. The DLMAZ swarm consortia potentially reflects the proposed transient existence of swarm consortia in nature. The only swarm consortia found to not include an *Archangium*, DLMAZ has no shared genetic features that may have passed horizontally between *Cystobacter* and *Microvirga*, and *C.* DLMAZ is not predicted to be a BCAA auxotroph. Adaptation of *M.* DLMAZ in a previous swarm consortia and loss of myxobacterial companion provides a potential explanation for discrepancies between it and other consortia. However, without further knowledge of the features involved, myxobacterial conditioning of symbionts and their movement between myxobacteria symbionts remains hypothetical.

Our findings demonstrate that *A*. WIMLSP2 exhibits markedly reduced predation toward its companion *M*. WIMLSP2, while retaining robust predatory capacity against non-symbiotic prey. Importantly, this attenuation is not attributable to a generalized defect in killing, as both *A*. WIMLSP2 and the model myxobacterium *My. xanthus* DK1622 displayed comparable predation efficiencies against *E. coli* (28% and 30%, respectively). Instead, the reduced susceptibility of *Microvirga* appears specific to its interaction with *Archangium*, consistent with an active, partner-specific modulation of predatory behavior. Although the molecular basis of this protection remains unresolved, myxobacteria are known to rely on cell-surface recognition systems to regulate social interactions, including kin discrimination and cooperative behaviors (3, 5). One possibility is that *Microvirga* presents surface features that are recognized by *Archangium*, thereby dampening the initiation or execution of predation. Alternatively, *Microvirga* may produce inhibitory factors, such as anti-toxins or signaling molecules, which interfere with downstream killing mechanisms following contact. Distinguishing between recognition-based avoidance and biochemical inhibition will be essential for defining how this interaction is regulated. More broadly, these findings suggest that myxobacterial predation can be selectively tuned toward particular neighbors, providing a potential mechanism by which stable associations and cooperative behaviors emerge within otherwise antagonistic microbial communities.

Previously described proto-farming of bacterial symbionts by the predatory, social amoeba *Dictystelium discoideum* serves as an established example of farming symbiosis between microbial predators and prey (85–87). Brock et al. characterized a farming phenotype present in natural isolates of *D. discoideum* that demonstrated bacterial husbandry (85). Farming *D. discoideum* clones curtail predation of bacteria and utilize fruiting body formation and subsequent sporulation to carry and seed bacterial crops during spore dispersal, and access to the dispersed food source in the absence of bacterial prey benefits farmers (85–87). Access to *Microvirgal* cells for nutrition in the absence of suitable prey would similarly benefit myxobacteria in swarm communities. Other parallels between farming symbiosis in *D. discoideum* and myxobacterial swarm consortia include the participation of social, cooperative predators with reduced predation of symbionts. Access to cultivable swarm consortia provides the opportunity to further investigate potential symbiotic farming by myxobacteria.

Myxococcota are a ubiquitous, keystone taxon in soil community structure with an outsized impact on nutrient cycling in the soil food web. Our results afford foundational insight into inter-bacterial symbiosis attributable to phenotypic selection of predation-resistant populations in polymicrobial communities. Although we have not determined if any swarm consortia members are obligate symbionts, we suspect this is a potential outcome in nature that may contribute to discrepancies between monocultured myxobacteria and myxobacteria present in metagenomic analysis of soil. Continued investigation of swarm consortia and comparison of xenic and axenic myxobacterial cultures will expand the current understanding of myxobacterial symbiosis, development, metabolism, and predation in polymicrobial communities.

## Methods

### Isolation and cultivation of swarm consortia

All swarm consortia were isolated from rhizospheric soils samples using methodology previously described for isolating myxobacteria from soil (20). Briefly, a lawn of *E. coli* was cultivated overnight at 37 °C on Luria–Bertani (LB) agar (1.5%). The resulting biomass was scraped into 2 mL of antifungal solution containing 250 µg/mL cycloheximide and nystatin. Approximately 300 µL of this suspension was then spread onto the center of WAT agar plates to generate a bait circle roughly two inches in diameter. Plates were allowed to air-dry to form *E. coli*–WAT bait plates. Separately, previously air-dried soil was rehydrated with the same antifungal solution to a mud-like consistency. Once the *E. coli* lawn on the WAT plate had dried, a pea-sized portion of the soil paste was placed at the center of the bait circle. Inoculated plates were incubated at 25 °C for up to four weeks and monitored daily for the development of lytic zones or fruiting bodies within the *E. coli* lawn. Emerging lytic zones were transferred using a sterile syringe needle onto VY/4 agar (2.5 g/L Baker’s yeast, 1.36 g/L CaCl_2_·2H_2_O, 0.5 mg/L vitamin B_12_, 15 g/L agar). The advancing swarm edge was repeatedly subcultured onto fresh VY/4 plates to obtain isolated colonies. However, throughout this purification process, the four samples, WIMLSP1, WIMLSP2, DLMAZ and FLWO consistently formed xenic swarms and remained resistant to monoculture despite multiple rounds of passaging. All swarm consortia were maintained on VY/4 plates and liquid cultures with CYH/2 media (0.75 g/L of casitone, 0.75 g/L of yeast extract, 2 g/L of starch, 0.5 g/L of soy flour, 0.5 g/L of glucose, 0.5 g/L of MgSO_4_•7H_2_O, 1 g/L of CaCl_2_•2H_2_O, 6 g/L of HEPES, 8 mg/L of EDTA-Fe, and 0.5 mg/L of vitamin B_12_). WIMLSP1 and WIMLSP2 were isolated from soil collected in Spring 2021 from the roots of a white spruce tree near White Lake, Michigan, USA (42.35, −84.35). FLWO was isolated from soil collected in Spring 2022 from the roots of a white oak tree near Palm Coast, Florida, USA (29.53, −81.22). DLMAZ was isolated from soil collected in Spring 2021 from the roots of a New Mexico desert locust tree near Mesa, Arizona, USA (33.41, −111.83).

### Scanning electron microscopy

Bacterial samples intended for scanning electron microscopy (SEM) were prepared using a standard fixation-dehydration protocol optimized for microbial colonies (82). Briefly, cells from each swarm consortia or monoculture myxobacterium after 72 h of growth on VY/4 agar were fixed in 2.5% glutaraldehyde prepared in 0.1 M phosphate buffer (pH 7.2) for 45 minutes to 1 hours, followed by 15 minutes wash in the same buffer. Samples were then re-fixed in 1% osmium tetroxide in PBS buffer for 60 minutes at room temperature to enhance membrane contrast and subsequently rinsed with buffer. Fixed samples were dehydrated through a graded ethanol series (30%, 50%, 70%, 90%, 95%, and 100%, in 10 min each), and the samples were dried overnight in a flow hood. Dehydrated samples were mounted onto aluminum stubs using carbon adhesive tape. Mounted samples were sputter-coated with a 10-20 nm layer of gold-palladium alloy using a Denton Vacuum Desk V TSC Sputter Coater and subsequently examined using a JEOL JSM-7200FLV Field-Emission Scanning Electron Microscope in the Glycore Imaging and Microscopy Core for SEM analysis.

### Metagenomic sequencing

High–molecular-weight metagenomic DNA was extracted from xenic swarm consortia using Qiagen Genomic-Tip columns. DNA concentration and purity were assessed using Qubit® dsDNA HS Assay Kits (ThermoFisher Scientific) and a Nanodrop spectrophotometer. Sequencing libraries were prepared according to the manufacturer’s instructions for the Native Barcoding Kit 24 V14 (Oxford Nanopore). The barcoded library was then loaded onto an Oxford Nanopore MinION flow cell (R10.4.1) and sequenced until approximately 1–2 Gbases of data per barcode were obtained. Raw nanopore reads were basecalled and demultiplexed with Dorado (v0.7.2) using the super-accurate model. Metagenome assembly was carried out with Flye (v2.9.2+), and consensus polishing was performed using Medaka (v1.11+) (83, 84). Resulting assemblies were deposited at the Joint Genome Institute Integrated Microbial Genomes & Microbiomes (JGI IMG/MER) database.

### Comparative genomic analysis

Annotation of sequenced swarm consortia metagenomes was completed using the JGI IMG annotation pipeline (v5.2.1) (44). Individual MAGs from swarm consortia were analyzed at the (TYGS) to acquire dDDH and 16S rDNA gene sequence comparisons (85–87). Average nucleotide identity values were calculated using the OrthoANI Tool (OAT) (v0.93.1)(88). Genome-scale metabolic models were constructed and characterized from MAGs using the MS2 Build metabolic models with OMEGGA (v2.0) and Run Model Characterization (v2.2.1) applications in KBase(53–55). Amino acid auxotrophies for each MAG were also calculated using GapMind(56). KEGG modules, KEGG pathways, and gene phylogenies associated with each metagenome were provided as part of the JGI IMG annotation pipeline and were accessed from the JGI IMG/MER database(44, 58–60). MEGA X (v10.1.7) was used for alignments of acyl-homoserine lactone synthases and ANKYR proteins and construction of phylogenetic trees (89). Predicted structures for acyl-homoserine lactone synthases and ANKYR proteins were modelled using the AlphaFold Server (AlphaFold 3) and subsequently aligned using the FoldMason Multiple Protein Structure Alignment tool (FoldMason MSA) (48, 49). ANKYR protein sequences from WIMLSP1 and WIMLSP2 were analyzed using the Enzyme Function Initiative Enzyme Functionality Tool (ESI-EST) to generate sequence similarity networks and determine phylogenetic distribution of homologs (50, 51, 90).

### Predation assays

WIMLSP2 was cultured in CY/H medium at 33 °C with shaking at 285 rpm overnight to approximately 200-300 Klett units to ensure each member was represented sufficiently. *M. xanthus* DK1622-GFP was grown in CTT broth (1% Casitone, 10 mM Tris-HCl pH 7.6, 8 mM MgSO₄, 1 mM KH₂PO₄) at 33 °C with shaking at 300 rpm to 80–120 Klett units. *E. coli* DH5α-mCherry was grown in LB supplemented with kanamycin (50 µg mL⁻¹) at 37 °C with shaking at 250 rpm to an OD₆₀₀ of ∼0.5.

For microscopy assays, cultures were back-diluted to 50 Klett units for the WIMLSP2 consortium and DK1622-GFP or to an OD₆₀₀ of 0.05 for *E. coli*. Three conditions were prepared for imaging: the WIMLSP2 consortium alone, WIMLSP2 mixed with DK1622-GFP, and WIMLSP2 mixed with *E. coli*-mCherry. Mixed samples were combined at a 1:1 (vol/vol) ratio immediately before imaging. A 5 µL aliquot of each culture or culture mixture was spotted onto TPM agar pads (10 mM Tris-HCl pH 7.6, 8 mM MgSO₄, 1 mM KH₂PO₄, 1% agar) supplemented with 2 mM CaCl₂. Pads were cast onto glass microscope slides and allowed to air dry before imaging.

Time-lapse fluorescence microscopy was performed using an Olympus IX83 inverted microscope equipped with a 60× oil-immersion objective and controlled with cellSens software. GFP and mCherry fluorescence were visualized using standard FITC and mCherry filter sets, respectively. Images were acquired every 20 seconds for a total duration of 2 hours. Predator–prey behaviors were scored manually from time-lapse movies. An interaction event was defined as the first frame in which a predator cell made contact with a prey cell. If the contacted prey cell underwent visible lysis within ten minutes, the event was scored as a kill; if no lysis occurred, it was scored as a non-lethal interaction. Repeated contacts between the same predator and prey cell were counted as separate interactions only when the cells were clearly separated and then re-established contact.

## Acknowledgements

This research was supported by funds from the National Institute of General Medical Sciences (R01GM149795 to D.C.S. and R35GM140886 to D.W.). We also acknowledge and appreciate access to the Glycoscience Center of Research Excellence Imaging Research Core and SEM assistance from John Adams Sabestian (P20GM130460).

Isolation, cultivation, and sequencing of swarm consortia: A.A., N.S., S.K.P., and B.T.; comparative genomics and bioinformatics: S.K.P. and D.C.S.; predation assays: S.W., D.B., and B.L.; conceptualization, manuscript preparation, and editing: N.S., S.K.P., D.C.S., D.W., and S.W.; supervision and administration: D.C.S. and D.W. All authors have read and approved the final manuscript.

## Supplemental Material

Supplemental data includes the following: Metagenome assembly statistics for swarm consortia (Tables S1-S4), swarm consortia details from assembly data (Table S5), dDDH data for swarm consortia myxobacteria and *Microvirga* (Tables S6 and S7), genes that are phylogenetically associated with Myxococcota that are present in *Microvirga* MAGs (Table S8), Genome BLAST Distance Phylogeny (GBDP) tree generated from myxobacterial 16S rDNA gene sequences (Figure S1), GBDP tree generated from myxobacterial MAG sequences (Figure S2), GBDP tree generated from *Microvirga* 16S rDNA gene sequences (Figure S3), GBDP tree generated from *Microvirga* MAG sequences (Figure S4), Phylogenetic distribution of ANKYR proteins identified with EFI-EST analysis (Figure S5), Alignment of genes encoding ANKYR proteins from WIMLSP1 and WIMLSP2 (Figure S6), Conserved spatial organization of XerC and ANKYR proteins in genomes of *M.* WIMLSP1, *M.* WIMLSP2, *M. solisilvae*, and *M.* sp. ACRRW (Figure S7), BCAA biosynthetic pathways from swarm consortia depicting BCAA auxotrophy in *Archangium* (Figure S8), Time-lapse of *A.* WIMSLP2/*M.* WIMSLP2 on TPM 1% agar pad after spot dried (Movie S1), Time-lapse of WIMLSP2/*E. coli-*mCherry on TPM 1% agar pad after spot dried (Movie S2), Time-lapse of WIMLSP2/DK1622-GFP on TPM 1% agar pad after spot dried (Movie S3), and Time-lapse of DK1622-GFP/ *E. coli-*mCherry on TPM 1% agar pad after spot dried (Movie S4).

## Data Availability

The data sets presented in this study can be found in online repositories. JGI Genomes Online Database (GOLD) Analysis Project IDs are as follows: WIMLSP1 (Ga0669685), WIMLSP2 (Ga0646599), FLWO (Ga0654980), and DLMAZ (Ga0646598).

## Notes

### Competing Interest Statement

The authors have declared no competing interest.

