## Supplemental movie legends for "Predator avoidance promotes inter-bacterial symbiosis with myxobacteria in polymicrobial communities"

**Supplemental movies**

**Supplemental movie S1.** Time-lapse of *A.* WIMSLP2/*M.* WIMSLP2 on TPM 1% agar pad after spot dried. Frame intervals 20 s over two hours with 60x objective. Arrows indicates lyses events of *Microvirga* (small rods) by *Archangium* (long rods). Some *Archangium* cells can be seen moving out of focus.

**Supplemental movie S2.** Time-lapse of WIMLSP2/*E. coli*-mCherry on TPM 1% agar pad after spot dried. Frame intervals 20 s over two hours with 60x objective. Arrows indicates lyses events of *E. coli*-mCherry (red) by *Archangium* (long rods).

**Supplemental movie S3.** Time-lapse of WIMLSP2/DK1622-GFP on TPM 1% agar pad after spot dried. Frame intervals 20 s over two hours with 60x objective. Arrows indicates lyses events of *Microvirga* (small rods) by DK1622-GFP (green).

**Supplemental movie S4.** Time-lapse of DK1622-GFP/ *E. coli*-mCherry on TPM 1% agar pad after spot dried. Frame intervals 20 s over two hours with 60x objective. Arrows indicates lyses events of *E. coli*-mCherry (red) by DK1622-GFP (green).
