## Supplemental Tables and Figures for "Predator avoidance promotes inter-bacterial symbiosis with myxobacteria in polymicrobial communities"

**Supplemental Data**

**Supplemental Table S1:** Assembly statistics for WIMLSP1

| Scaffold ID (IMG/MER) | Sequence length (bp) | GC content | Gene count |
|---|---|---|---|
| Ga0669685_01 | 12,981,101 | 0.69 | 10,693 |
| Ga0669685_02 | 2,513 | 0.64 | 6 |
| Ga0669685_03 | 3,923,152 | 0.61 | 3,882 |
| Ga0669685_04 | 2,538 | 0.68 | 2 |
| Ga0669685_05 | 2,575 | 0.70 | 3 |
| Ga0669685_06 | 2,776 | 0.69 | 2 |

**Supplemental Table S2:** Assembly statistics for WIMLSP2

| Scaffold ID (IMG/MER) | Sequence length (bp) | GC content | Gene count |
|---|---|---|---|
| Ga0646599_01 | 12,928,281 | 0.69 | 10,576 |
| Ga0646599_02 | 4,285,839 | 0.62 | 4,146 |
| Ga0646599_03 | 5,931 | 0.56 | 13 |

**Supplemental Table S3:** Assembly statistics for FLWO

| Scaffold ID (IMG/MER) | Sequence length (bp) | GC content | Gene count |
|---|---|---|---|
| Ga0654980_01 | 13,960,269 | 0.68 | 11,322 |
| Ga0654980_02 | 2,321 | 0.65 | 2 |
| Ga0654980_03 | 2,877 | 0.69 | 2 |
| Ga0654980_04 | 1,081 | 0.66 | 3 |
| Ga0654980_05 | 1,603 | 0.63 | 2 |
| Ga0654980_06 | 1,183 | 0.64 | 1 |
| Ga0654980_07 | 3,818,472 | 0.61 | 3,738 |
| Ga0654980_08 | 1,347 | 0.67 | 1 |

| | | | |
|---|---|---|---|
| Ga0654980_09 | 5,895 | 0.56 | 14 |

**Supplemental Table S4:** Assembly statistics for DLMAZ

| Scaffold ID (IMG/MER) | Sequence length (bp) | GC content | Gene count |
|---|---|---|---|
| Ga0646598_01 | 44,387 | 0.58 | 50 |
| Ga0646598_02 | 12,010,403 | 0.69 | 10,262 |
| Ga0646598_03 | 19,741 | 0.71 | 23 |
| Ga0646598_04 | 158,443 | 0.68 | 86 |
| Ga0646598_05 | 4,194 | 0.68 | 7 |
| Ga0646598_06 | 12,788 | 0.72 | 13 |
| Ga0646598_07 | 133,616 | 0.59 | 113 |
| Ga0646598_08 | 7,740 | 0.72 | 12 |
| Ga0646598_09 | 6,295 | 0.70 | 9 |
| Ga0646598_10 | 4,131,692 | 0.61 | 4,039 |

**Supplemental Table S5:** Swarm consortia details from assembled data

| Swarm consortia | contigs | # of bases | # of RNA genes | # of protein coding sequences | Estimated # of genomes |
|---|---|---|---|---|---|
| WIMLSP1 | 6 | 16,914,655 | 182 | 14,387 | 2 |
| WIMLSP2 | 3 | 17,220,051 | 181 | 14,534 | 2 |
| FLWO | 9 | 17,795,048 | 180 | 14,889 | 2 |
| DLMAZ | 10 | 16,529,299 | 156 | 14,442 | 2 |

**Supplemental Table S6:** dDDH data for swarm consortia myxobacteria. dDDH ($d_4$, in %) values provided by TYGS analysis of corresponding MAGs. dDDH values were not calculated for cells with “n.d.”

| | A. WIMLSP1 | A. WIMLSP2 | A.FLWO | C.DLMAZ | A.gephyra | A.lansingense | C.fuscus |
|---|---|---|---|---|---|---|---|
| **A.WIMLSP1** | 100 | 92.7 | 34.8 | 25.3 | 45.9 | 36.2 | 24.9 |
| **A.WIMLSP2** | 92.7 | 100 | 34.8 | 25.1 | 45.7 | 36.2 | 24.9 |
| **A.FLWO** | 34.8 | 34.8 | 100 | 25.2 | 35.4 | 44.9 | 24.9 |
| **C.DLMAZ** | 25.3 | 25.1 | 25.2 | 100 | 25.2 | 25.9 | 52.3 |
| **A.gephyra** | 45.9 | 45.7 | 35.4 | 25.2 | 100 | n.d. | n.d. |
| **A.lansingense** | 36.2 | 36.1 | 44.9 | 25.9 | n.d. | 100 | n.d. |
| **C.fuscus** | 24.9 | 24.9 | 24.9 | 52.3 | n.d. | n.d. | 100 |

**Supplemental Table S7:** dDDH data for swarm consortia *Microvirga*. dDDH ($d_4$, in %) values provided by TYGS analysis of corresponding MAGs. dDDH values were not calculated for cells with “n.d.”

| | M. WIMLSP1 | M. WIMLSP2 | M.FLWO | M.DLMAZ | M.guangxiensis | M.solisilvae | M.vignae |
|---|---|---|---|---|---|---|---|
| **M.WIMLSP1** | 100 | 25.9 | 38 | 25.9 | 30.3 | 28.5 | 27.8 |
| **M.WIMLSP2** | 25.9 | 100 | 25.6 | 38.5 | 24.9 | 25.3 | 24.8 |
| **M.FLWO** | 38 | 25.6 | 100 | 25.8 | 30.2 | 28.5 | 27.7 |
| **M.DLMAZ** | 25.9 | 38.5 | 25.8 | 100 | 25.2 | 25.5 | 24.9 |
| **M.guangxiensis** | 30.3 | 24.9 | 30.2 | 25.2 | 100 | n.d. | n.d. |
| **M.solisilvae** | 28.5 | 25.3 | 28.5 | 25.5 | n.d. | 100 | n.d. |
| **M.vignae** | 27.8 | 24.8 | 27.7 | 24.9 | n.d. | n.d. | 100 |

**Supplemental Table S8:** Genes that are phylogenetically associated with Myxococcota that are present in *Microvirga* MAGs.

| MAG | AA sequence | annotation | Top BLASTP hit (species | 2nd BLASTP hit (species) |
|---|---|---|---|---|
| **M.WIMLSP1** | MLKLAFAAAVGTTLLFASGAQALETQNGLAFNGLA FNGLAFNGLAFNGLAFNGAAADGVAGELRSAPALQ ATTVILKDGERVSLK | hypothetical protein | Microvirga sp. ACRRW | Pyxidicoccus sp. 3LG |
| | MLMNQDEYDKHLKGFMITGCTVRSKDVFYLVAITD SPNRARPESDLTTRVIPYFFEKIEKRWGHINYHGYS RTLAGASLYPESKFVGVDRGGQVMVVGGGKMEIE DIAGGRTGPIRGSVNRVRTINGFIHVCSNNRGLARR DGTDRWTSLCADLPVKPNPNGFGEVYGFNDFDAF DNGEFYCVGGQSDVWRFDGANWTPIDVPGDHNG ASNFLTKAGSKIARVPLHSVCCAGDGYLYIGGPDG GVWKGRNEQWKLIHDDRLSLPFKDIVWFQDRVYC TSRYGLWEIVNDEVRPCDVPEEISICSGNLAVADGI MLLAGECGAAYHDGREWKLIFNTSSFT | WD40/YVTN/BNR-like beta propeller repeat protein | Microvirga sp. ACRRW | Archangium sp. |
| | MIFRHLRGGLRAVLFTLLGLLGYPSLVSAAGPDMYF DAPADKQMALAVASGDIETMTALLSSKAVDPQAIG RKATSWIEIAIIADQKKAFDTLMKWNALGPPKGKIA GQAMYSATVKGSIRWLERLAAAGASLDNYGGGEL LIVTALDTRNEAVLDFYIRNGADLDMPAMAGGSVAL SAAMTRRFDMALRFLDLGASPWVMDSLGSTLGSIA ERAARVPAWDHSSRMNQHRLELLQRLHAIGFPDP APTADEGHALRQKKQWPPKAAIKQ | ANKYR superfamily | Microvirga sp. ACRRW | Archangium sp. |
| | MFKIPEPMLTYWASKAYKKASIADLAEVERLFGTAL PASYVEFVTTIGFVVFDDVPGFKIHEYFDYKVGSPE GTEIAQGNIAFLKEPAHIIKAHKILTNRQALEEEEEED EDFPKFPKNYLPIANDAGQGQILMEFGEHPGRIWY WQENDWAWGLEDNTWLGFVAENFEDFINGLKP | SM1/KNR4 family protein | Archangium violaceum | Candidatus Methylumidphilus sp. |
| **M. WIMLSP2** | MSRTMKYTILVSFVGLLSAGAQAAEIANGSDLNGA NLNGANLNGSDLNGSDLNGANLNGASSGRVFLGA QVSALIAPDGTLVTLD | pentapeptide repeat protein | Microvirga sp. | Archangium violaceum |
| | MMTTIDDIRRRDGFLTIEETLSLCDRNTIYDPYSTLIS RHARIGSGNILYPCTTIRCSQDSSCEVGDRNIFHSL TMIDACGGAISIGSGNTFGDGGFTAKADRPGAKITI GDRGRYASGASVYGVSHLGTGSQILGQISVIDCVL ADGEDFTHADPDERAAVLKGHGAARKLRIGVGEVI FGKSSFDQAGIQRQTDFHPKS | IpxD-like protein | Pyxidicoccus sp. 3LG | Stappiacea bacterium |
| | MSYPFYLLYGAYLASGLGDWILRIAVPLLIYQVTDSA LAMAGAYAVNYLPYLIVTPFGGVLADRVDRRRMLL VGDFFAAGLVVAIILANASGAAALLLYPLLFVLASTA AVYHPGFQSFIPSVVPPDKLARANSLFAAADNGLSL LGPAAAGGIVALLGPVQALYADALSFALSGLLILCIP SAMSAKAQVERKLQGILHDLREGFVYVWHNRILRA GAFLFFFVNFSYDIFYANFIFLLVGIFGLTAVDAGTVI SMTGVGALVGSLVAPKLMSRVSSGRLIVACTATAG GLILLLLFVDGALAVGMLWGGVCATQAVIRVAYFTL RQKIVPSNLLGRSVAVTRMISYAAVPLAALSGGWIV QQTGEIRMIVIISGSVMLLSALIAWFTSLGRTPQPAL QSSLAT | MFS transporter | Microvirga sp. | Polyangium aurulentum |
| | MILRQLRGGLRALLFMLLGLFGYPASASAAGPDMY FDSPADKQMALAVASGDIETMTALLSSKAVDPLAIG RKVTSWIEIAVIADQRAAFDALVKWGALGPAKGKIA GQAMYSATIKGSIRWLERLTAAGASLDNHGGGDLL IVQALDTRNEAVLDFYIRNGADLNMPAMAGGSVAL SAAMTRRFDMVLKFLDLGASPWVMDSLGSTLGSIA | ANKYR superfamily | Microvirga sp. ACRRW | Archangium sp. |

| | | | | |
|---|---|---|---|---|
| | ERAARVPAWDHSSRMNQHRLELLRRLHAIGFPDPAPTANEGHALRQKKQWPPKAAIKQ | | | |
| **M. FLWO** | MRHRISIGLACILCLMGAGCLDEREKAARTLVKTRACPDCDLTEIKLEAAQLQGAQLAGARLEKADMRKADLREADFSGAILFDTDLRGADLRGAAFREASMTGAQMQGANLEGVDLSGTTLNAIDLSGVNFQGANLRGAKLSEARLNGINGPYRRDPPAHFYAVPAGGADLRGADLSGADLSGAYLSKADLRDAKLAGANLRDAHLDQADLRGADINGADLKGAILDHATWIDGSICAEKSIGRCRRP | pentapeptide repeat protein | Corallococcus sp. | Corallococcus silvisoli |
| | MKASKWQLKQRWHEEAGALVLKRVQDILLADDTRKLPLGIPDILSGLPFRDEVASGRDFRGIELEGGLTSLDLSGCDFSYAKLTLNFIRCDLSEANFEEATLGGIIFDKATRANFRRAKMRHCSLVGLNAQDCCFDEAILSNASFEKACLQGSKFRNANCKGASFVSANLLGCDFQGANLNECPFQGVILDRSTNLRGASLVGLFYHEHRSIDGKLVLPKTDWRLATHDETTRTEA | pentapeptide repeat protein | Hyalangium gracile | Archangium sp. |
| | MLKTAFAATVGTTLLFGSGAQALETQNGLAFNGLAFNGLAFNGLAFNGLAFNGAATDGVAEELRSAPALQATTVILKDGEHVSLK | hypothetical protein | Microvirga sp. ACRRW | Pyxidicoccus sp. 3LG |
| **M. DLMAZ** | MACAIGLYHRVKARIPTIAVICTDANSCYRLAFARHGVAEAHVQSKAHTH | transposase | Corallococcus sp. AB045 | Alphaproteobacteria bacterium |
| | MTCHHCGSTAFRKNGHCAGVQRYVCHACHRSFSANGERFSKAVKAQALDM | transposase | Corallococcus sp. AB045 | Accumulibacter sp. |
| | MGVLERLVLTDSQWARIAPLIIGRPDQKGSTGRDNRMFVEGVLWIVRTGS | transposase | Microvirga sesbaniae | Corallococcus sp. AB045 |
| | MACHHCGSSAFCKNGHTRGVQRYRCHACHRSFSANGERFSKTVKAQALDMYLNNVGLRKIARFTGASPPAVLKWIKKAATALAAQLEQAKAQVHDELPDVIEMDEIYTFVQKNSSAPSYGLLILDGRAVLLRTSSATGA | transposase | Corallococcus sp. AB045 | Accumulibacter sp. |
| | MSSAIGLYRRVKQAVPAVALICTDANSCYRLAFERYRVPEAHVQSKAHTH | transposase | Corallococcus sp. AB045 | Mesorhizobium sp. 8 |
| | MLLDQTAYDAYFKGFILIDCVIRSKDIFYFVLVSDLKRTRSESDRKTRIVAHFLKSSDKPWRRADYEGFAKVFAGASQLPASKFVGVDRGAQVMLIGSGSLENEDIPAGKQGPIRGAIRKIKTINGYAHVCSGYRGFARRDGPNLWTSLVKNLNFMPDPDKDSGIYGFADFDAFNDRDFYCVGGHSDAWHFDGETWTQLDFPGDPSQIPESLIDPSTPGVPLEAVCCAGNGYVYIGGPGGTVWKGRKNSWTLIHRDSMSLPLRDMVWFKDRVYCTSDYGLWEIVDDQLRPCDIPDEIRVCSGHLSVCDGVMLLAGIYGAAYHDGNRWHLIFDTGQF | WD40/YVTN/BNR-like beta propeller repeat protein | Microvirga sp. ACRRW | Archangium sp. |

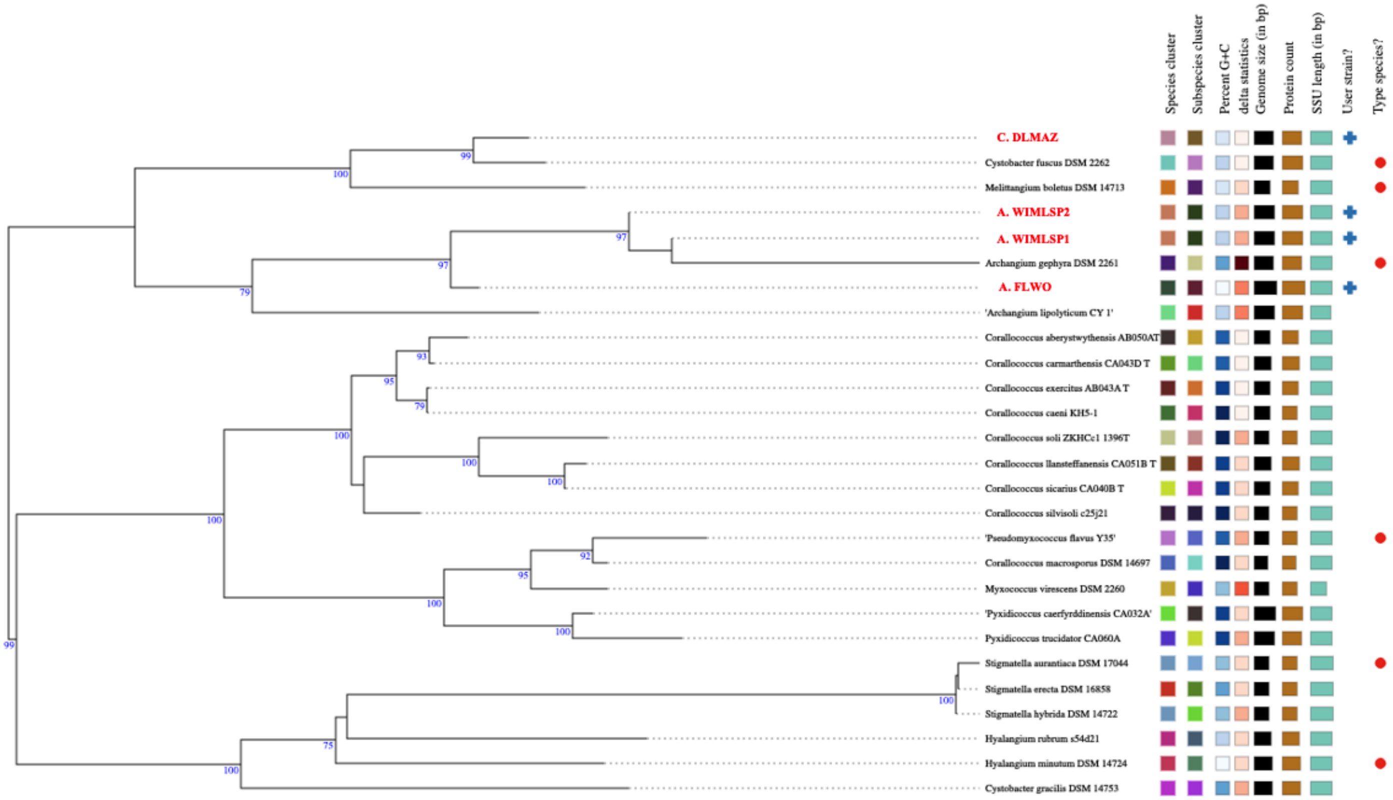

**Supplemental Figure S1:** Genome BLAST Distance Phylogeny (GBDP) tree generated from myxobacterial 16S rDNA gene sequences at the Type Strain Genome Server (TYGS).

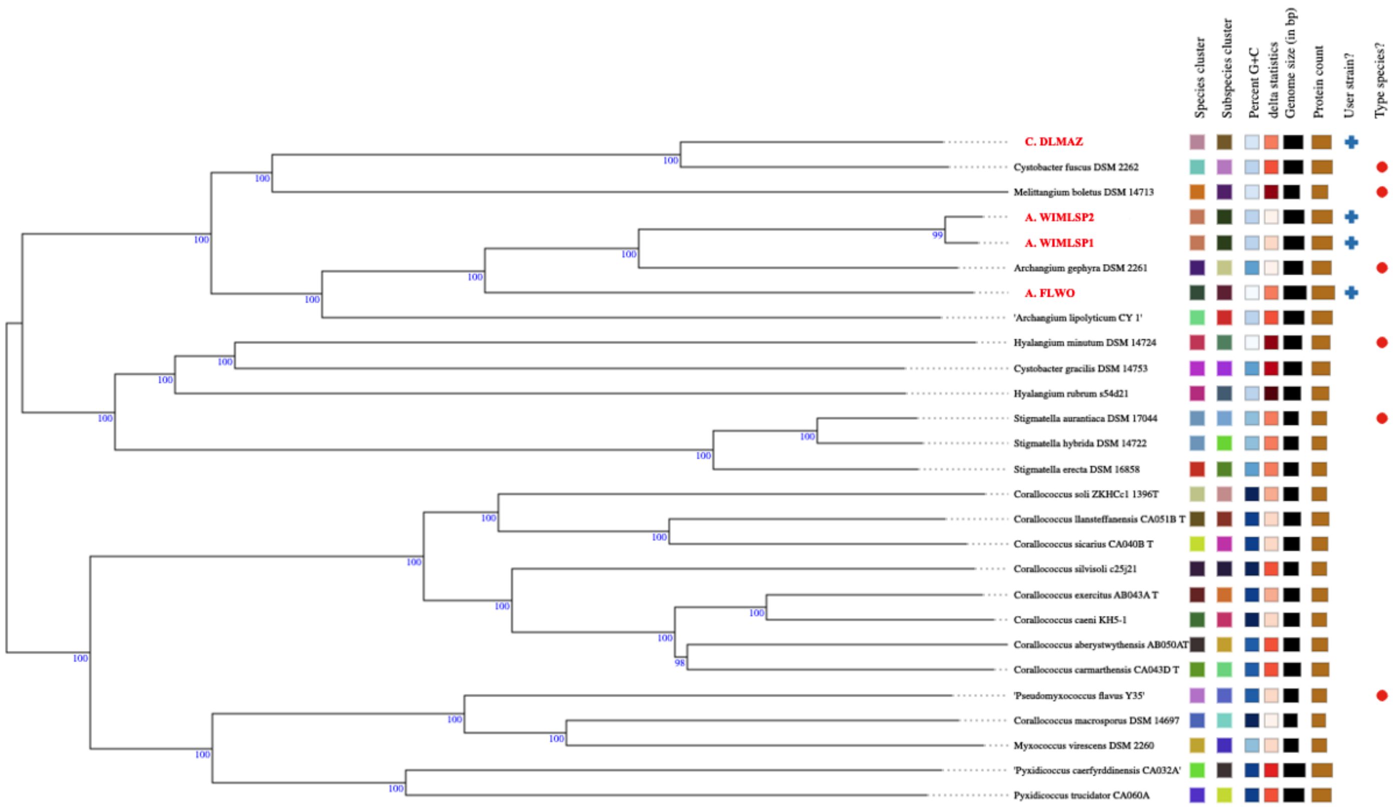

**Supplemental Figure S2:** Genome BLAST Distance Phylogeny (GBDP) tree generated from myxobacterial MAG sequences at the Type Strain Genome Server (TYGS).

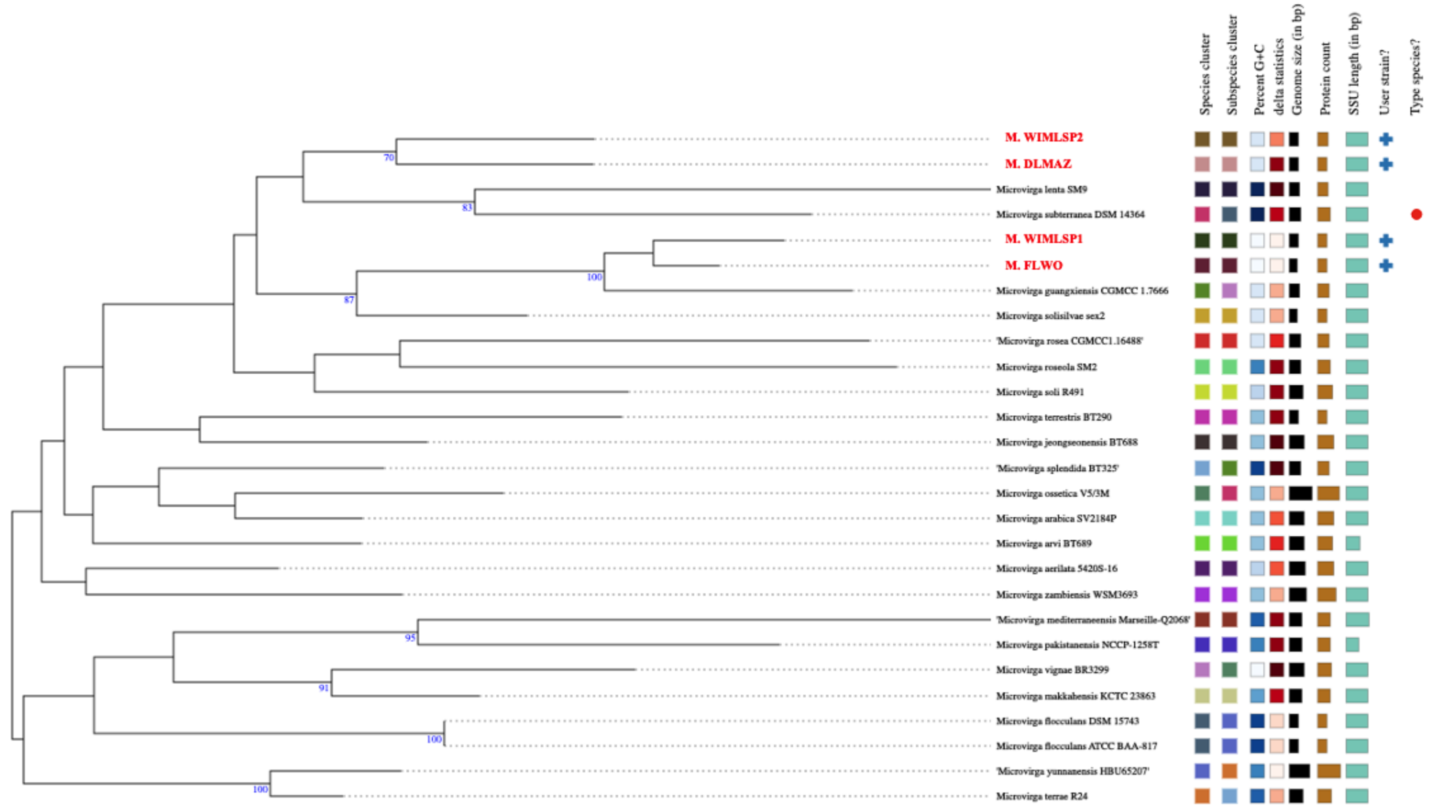

**Supplemental Figure S3:** Genome BLAST Distance Phylogeny (GBDP) tree generated from *Microvirga* 16S rDNA gene sequences at the Type Strain Genome Server (TYGS).

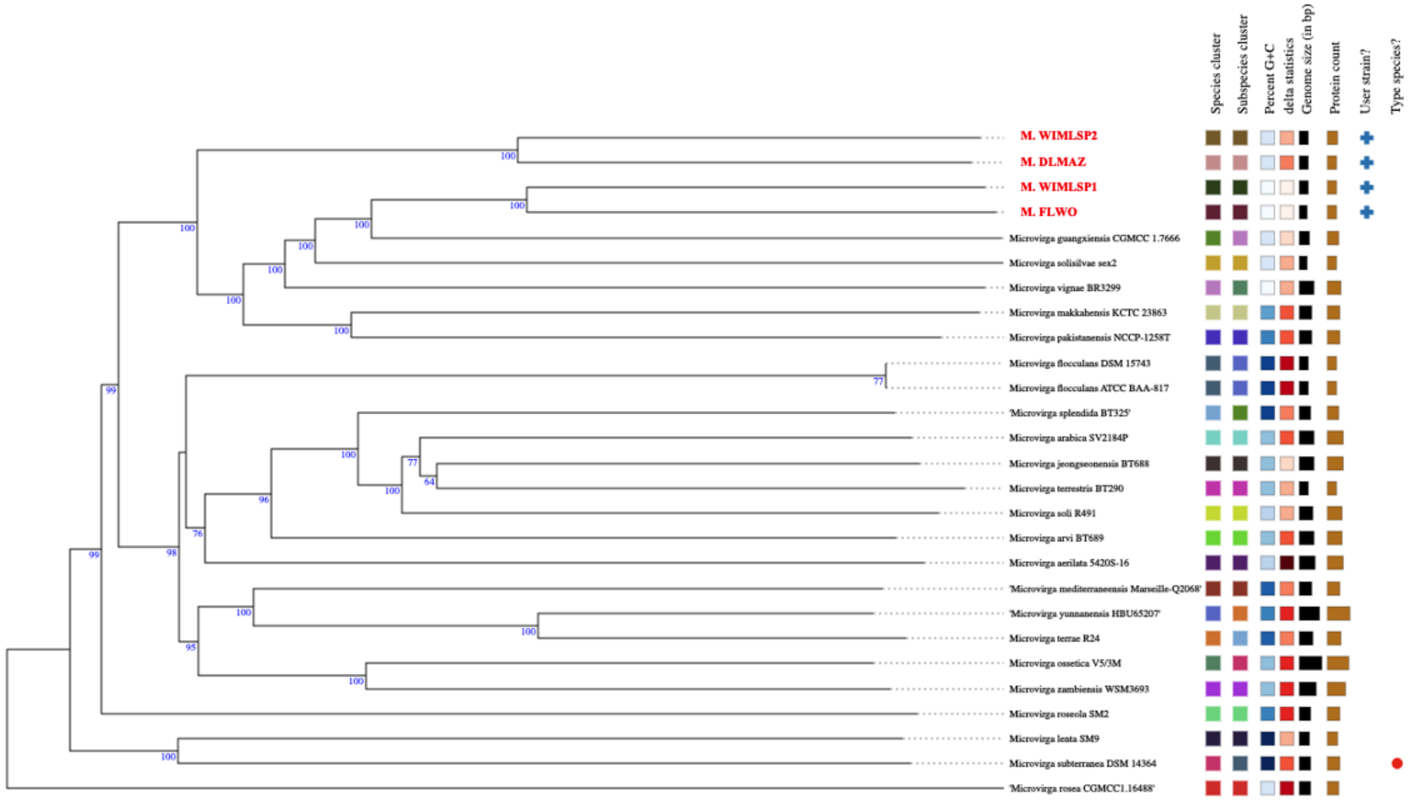

**Supplemental Figure S4:** Genome BLAST Distance Phylogeny (GBDP) tree generated from *Microvirga* MAG sequences at the Type Strain Genome Server (TYGS).

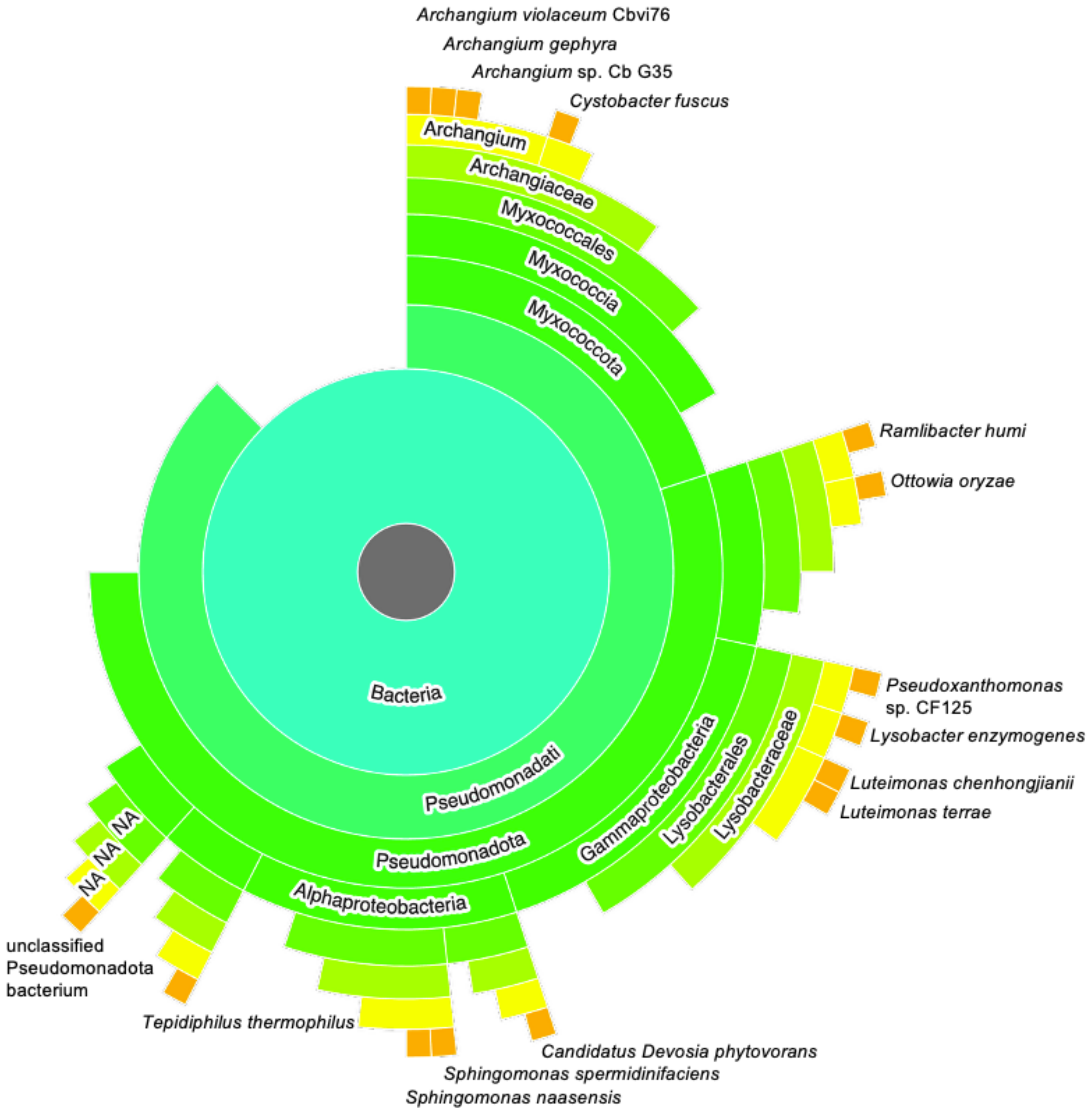

**Supplemental Figure S5:** Phylogenetic distribution of ANKYR proteins identified with EFI-EST analysis.

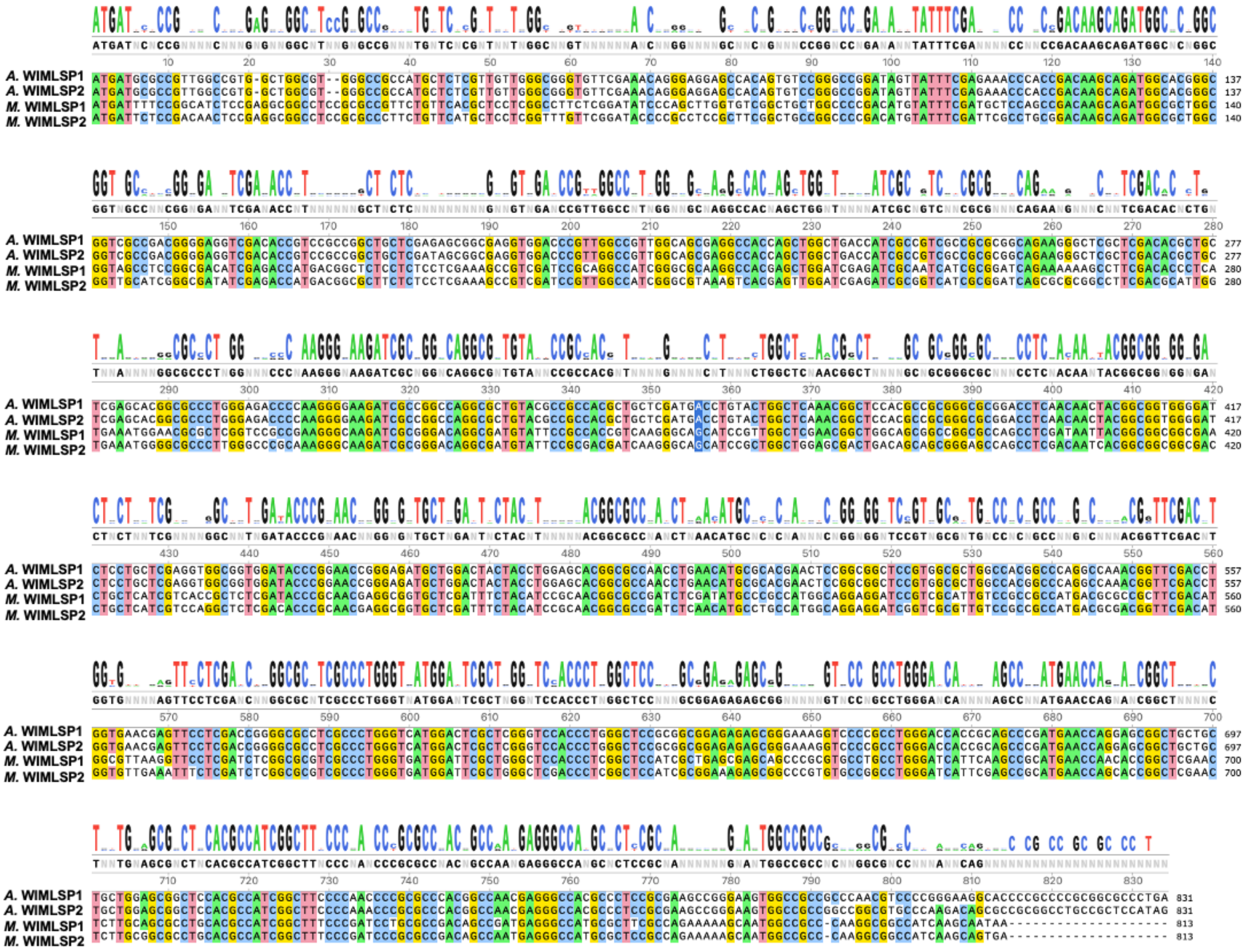

**Supplemental Figure S6:** Alignment of genes encoding ANKYR proteins from WIMLSP1 and WIMLSP2 generated with MEGA X using Clustal.

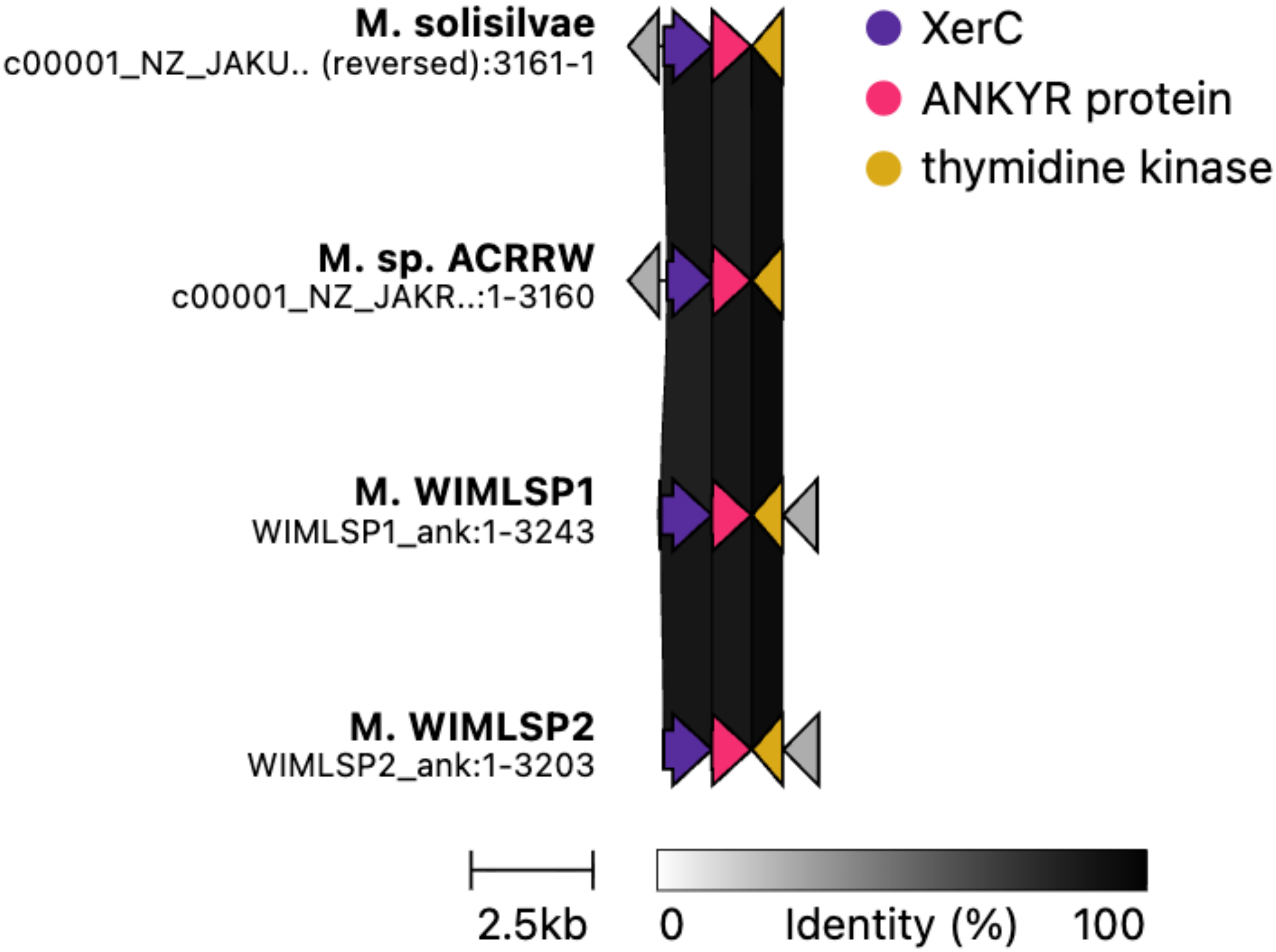

**Supplemental Figure S7:** Conserved spatial organization of XerC and ANKYR proteins in genomes of *M.* WIMLSP1, *M.* WIMLSP2, *M. solisilvae*, and *M.* sp. ACRRW. Figure rendered with clinker.

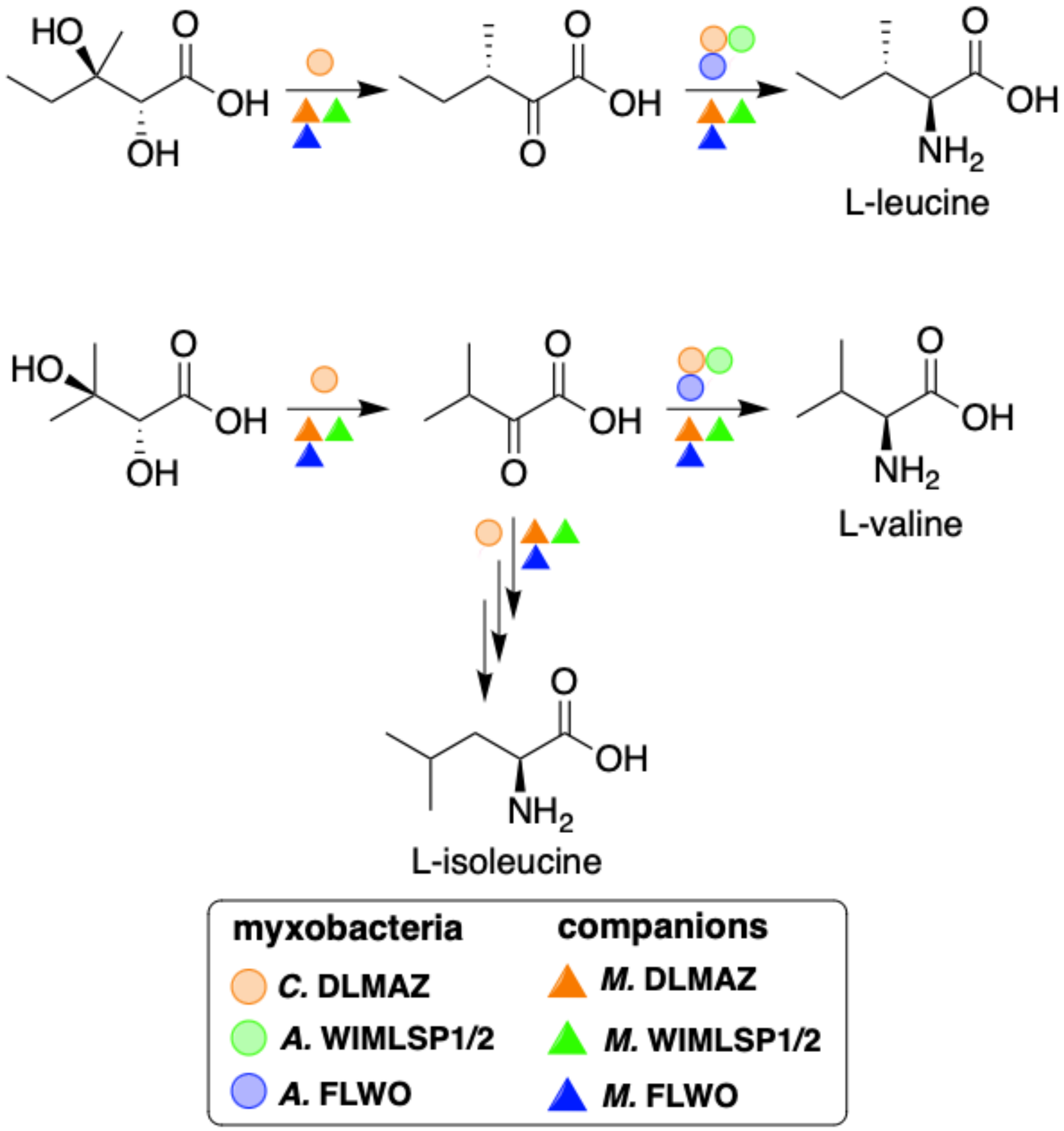

**Supplemental Figure S8:** BCAA biosynthetic pathways from swarm consortia depicting BCAA auxotrophy in *Archangium*.
